# The promiscuous development of an unconventional Qa1^b^-restricted T cell population

**DOI:** 10.1101/2022.09.26.509583

**Authors:** Michael Manoharan Valerio, Kathya Arana, Jian Guan, Shiao Wei Chan, Xiaokun Yang, Nadia Kurd, Angus Lee, Nilabh Shastri, Laurent Coscoy, Ellen A. Robey

## Abstract

MHC-E restricted CD8 T cells show promise in vaccine settings, but their development and specificity remain poorly understood. Here we focus on a CD8 T cell population reactive to a self-peptide (FL9) bound to mouse MHC-E (Qa-1^b^) that is presented in response to loss of the MHC I processing enzyme ERAAP, termed QFL T cells. We find that mature QFL thymocytes are predominantly CD8αβ+CD4-, show signs of agonist selection, and give rise to both CD8αα and CD8αβ intraepithelial lymphocytes (IEL), as well as memory phenotype CD8αβ T cells. QFL T cells require the MHC I subunit β-2 microglobulin (β2m), but do not require Qa1^b^ or classical MHC I for positive selection. However, QFL thymocytes do require Qa1^b^ for agonist selection and full functionality. Our data highlight the relaxed requirements for positive selection of an MHC-E restricted T cell population and suggest a CD8αβ+CD4-pathway for development of CD8αα IELs.

## Introduction

Thymic development of conventional CD8 T cells requires low affinity recognition of self-peptides bound to MHC I molecules expressed by cortical thymic epithelial cells and gives rise to naïve circulating CD8 T cells. Conventional CD8 T cells recognize peptides bound to classical MHC I (called MHC Ia) molecules, in contrast to unconventional T cell populations that recognize a diverse set of non-classical MHC I (called MHC Ib)^1,2^. MHC Ib molecules are structurally homologous to MHC Ia, and often associate with β2m, but are generally non-polymorphic, and can bind peptides or non-peptidic ligands ^3^. The two most prominent and well-studied examples of unconventional αβTCR-expressing T cells are mucosal associated invariant T cells (MAIT cells), that recognize vitamin B metabolites presented by MR1, and invariant natural killer T cells (iNKT cells), that recognize lipid metabolites presented by CD1d. MAIT cells and iNKT cells, like conventional T cells, require their cognate MHC ligand to develop in the thymus^4,5^. However, unlike conventional T cells, they undergo “agonist selection”, an alternative thymic selection process in which strong TCR signals do not lead to cell death, but instead drive alternative differentiation programs^6^. In addition, they recognize self-ligands presented by thymic antigen presenting cells (APCs) of hematopoietic, rather than epithelial, origin ^4,5^. The development of T cells restricted to other MHC Ib molecules remains understudied ^7–12^.

While MHC Ib restricted T cells are relatively rare in circulation, they contribute substantially to the intraepithelial lymphocyte (IEL) compartment of the small intestine ^13–15^. αβTCR+ IEL are generally classified as either induced or natural IELs, which differ in their specificity and developmental pathways. Induced IELs, which express the CD8αβ heterodimer, are specific for classical MHC Ia molecules and are derived from conventional CD8 T cells following antigen encounter in the periphery^16,17^. On the other hand, natural IELs, which predominantly express the CD8αα homodimer, can recognize a variety of different MHC ligands and are programmed for an IEL fate by strong recogniton of self ligands in the thymus^18,19^. Studies of natural IEL development have largely focused on populations of αβTCR+CD4-CD8- (double negative or DN) thymocytes, which can give rise to CD8αα IEL upon transfer into T cell deficient mice ^20,21^. However, it is unclear whether all natural IEL develop via an αβTCR+DN stage. Moreover, while it is known that many natural IEL require β2m, but not MHC Ia molecules, for their development^13–15,18,19^, the specifity of IEL for particular MHC Ib molecules remains largely unknown. As a result, no studies to date have focused on the development of IELs specific for defined MHC Ib molecules.

The MHC Ib molecule MHC-E (called Qa1 in mouse) is best known for its role in regulating NK cell responses; however, recent attention has focused on its function as a restricting MHC molecule for CD8 T cells^22,23^. In healthy cells, MHC-E molecules predominantly display a self-peptide derived from an MHC Ia leader peptide (called QDM peptide in mouse), which serves as a ligand for NK receptors and provides an inhibitory signal to NK cells and activated T cells ^24,25^. However, under conditions of impaired MHC Ia presentation, such as deficiency in ERAAP (endoplasmic reticulum aminopeptidase associated with antigen processing) or TAP (transporter associated with antigen processing), Qa1^b^ is loaded with an alternative set of peptides that can be recognized by CD8 T cells^10,26–28^. MHC-E restricted T cells responsive to TAP and ERAAP deficient cells have been proposed to play a role in monitoring defects in MHC Ia presentation induced by viral infection^29^ transformation or stress^27,28^. In addition, pathogen-specific MHC-E restricted CD8 T cells can be activated upon infection with a variety of viruses and bacteria ^30–34^. Recent studies of a CMV-vectored anti-HIV vaccine showed that MHC-E restricted CD8 T cells can produce responses that are extremely broad, with an unusually large proportion of the potential epitopes being targeted for recognition, and which provide strong immune protection^35,36^. Altogether, the ability of MHC-E restricted T cells to respond broadly to both microbial antigens and abnormal self, suggests an unusual mode of T cell recognition with significant therapeutic potential. However, our limited understanding of the specificity and development of MHC-E restricted CD8 T cells hampers our ability to harness these responses for therapeutic purposes.

Perhaps the best characterized example of an MHC-E restricted CD8 T cell response is QFL T cells, which recognize <u>Q</u>a1^b^ loaded with a self-peptide FYAEATPML (<u>FL</u>9) derived from Fam49a/b proteins^27^. QFL T cells were discovered as part of the mouse T cell response upon immunization of wild type mice with ERAAP deficient splenocytes. Interestingly, QFL T cells display hybrid characteristics of both conventional and unconventional T cells. Like conventional MHC Ia-restricted T cells, QFL T cells are found in the spleen and express the CD8αβ heterodimer. However, reminiscent of MAIT and iNKT cells, the majority use a semi-invariant TCR with a fixed TCRα and limited TCRβ usage ^37^. Splenic QFL T cells display an antigen experienced phenotype in wild type, unimmunized mice, reminiscent of conventional CD8 T cells that acquire a memory phenotype following homeostatic proliferation to self, termed “memory phenotype” or “virtual memory” T cells ^38,39^. While QFL T cells can be detected using FL9-Qa1^b^ tetramers (called QFL tetramers) in wild type and Qa1^b^ deficient mice^27^, their development in the thymus, and their contribution to the IEL compartment have not yet been examined.

Here we use both QFL tetramers and mice expressing rearranged QFL-specific αβTCR transgenes to probe the development of QFL T cells in wild type and MHC I deficient mice. QFL T cells can be readily detected in the spleen, thymus, and IEL compartment, with QFL T cells in the IEL compartment comprised of both CD8αα and CD8αβ phenotypes. Our data indicate that Qa1^b^ expression, predominantly by hematopoietic cells, drives the agonist selection of QFL T cells in the thymus, leading to mature CD8+CD4- thymocytes that exhibit signs of strong TCR signals. However, QFL T cells also recognize an alternative MHC I ligand, which can allow for positive selection of QFL CD8SP thymocytes with a more conventional phenotype in the absence of Qa1^b^. Our data highlight the promiscuous recognition and development of QFL T cells, confirm their hybrid conventional/unconventional characteristics, and suggest an alternative pathway for the development of natural IELs.

## Results

### Characterization of QFL T cells in wild type and TCR transgenic mice

To investigate the development of QFL specific T cells, we used tetramer enrichment of lymphocytes using Qa1^b^ tetramers loaded with the FL9 peptide^27^(hereafter called QFL tetramers). To increase the specificity of detection, we co-stained using both the QFL tetramer and an antibody specific for Vα3.2, which recognizes the invariant TCRα chain used by the majority of QFL T cells (Supplementary Fig. 1A-B) (Fig. 1A) ^37^. In this study, we focused on QFL tetramer^+^ and Vα3.2^+^ cells, hereafter called QFL T cells. The majority of QFL T cells in the thymus, spleen and small intestine (SI) intraepithelial lymphocyte (IEL) compartment of wild type mice were CD8α^+^CD4- (Fig. 1A). Interestingly, while mature QFL T cells in thymus and spleen predominantly expressed the CD8αβ heterodimer, QFL T cells in the IEL compartment were a mixture of cells expressing CD8αα or CD8αβ (Supplementary Fig. 1D-E). As previously reported^27^, QFL T cells were relatively abundant in the spleen of wild type mice (∼ 1/6197 of CD8 T cells or ∼1325/spleen/mouse, Fig. 1B-C). For comparison, a study of conventional CD8 T cell frequencies reported a range of 1/30,000 to 1/160,000 ^40^. Additionally, a substantial number of QFL T cells were identified in the thymus and IEL compartment of the small intestine, with an average of 245 QFL T cells and 1,174 QFL T cells respectively (Fig. 1B). The frequency of QFL T cells out of mature CD8 T cells was higher in the SI IEL compared to the thymus and spleen (Fig. 1C), suggesting that they undergo selective recruitment and/or expansion in this compartment.

**Figure 1:**
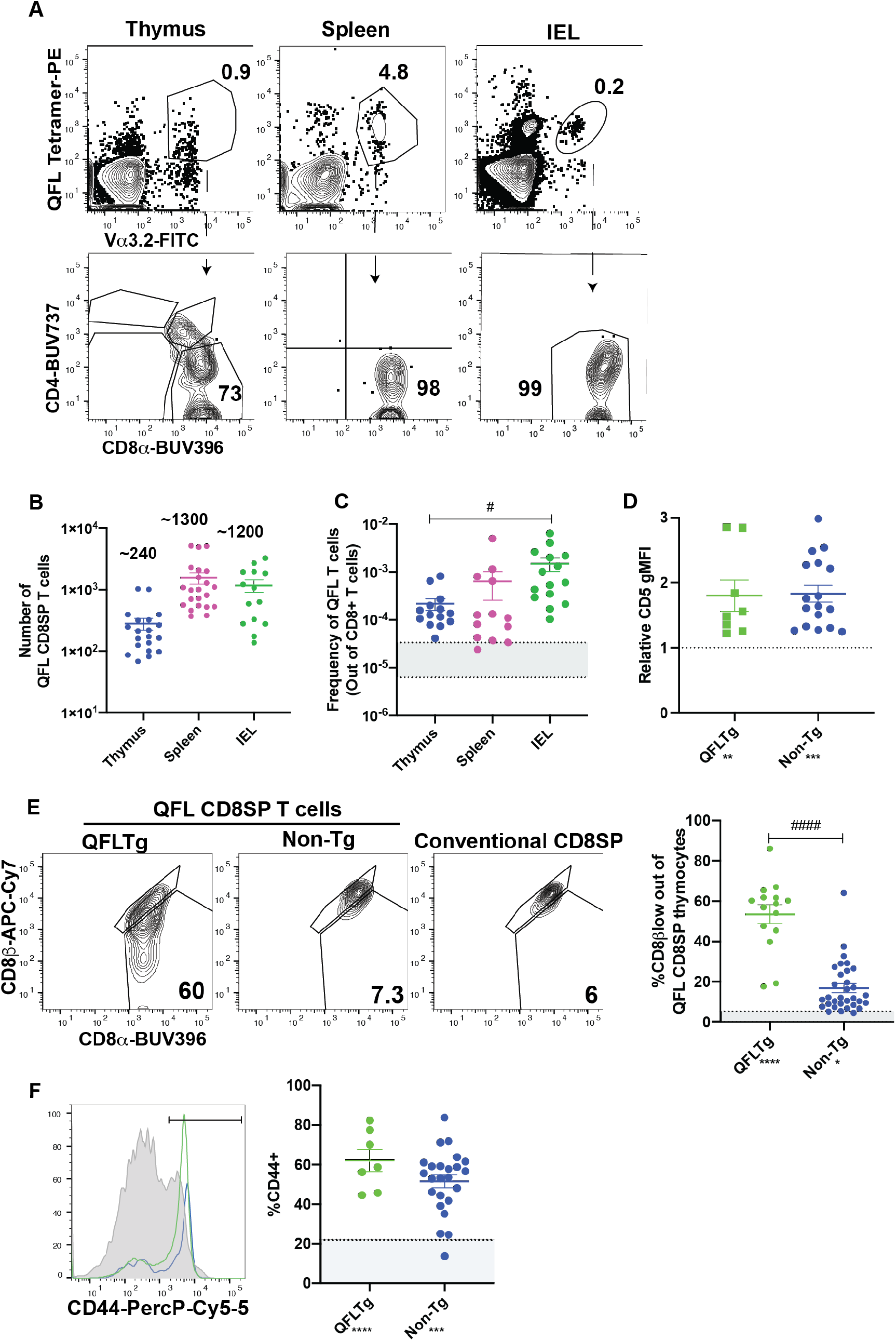
Characterization of QFL T cells in non-transgenic and transgenic mice. **A**-**C)** QFL T cells were identified by flow cytometry from wild type mice. **A)** Representative plots of QFL tetramer and Vα3.2 TCR from tetramer enriched thymocytes, tetramer enriched splenocytes, and unenriched small intestine intraepithelial lymphocytes. Splenocytes and IEL were gated for TCRβ+ cells. CD4 and CD8α expression on the indicated gated populations are shown below. **B)** Absolute numbers of QFL CD8SP T cells in the indicated compartments of wild type mice (Thymus n=14, IEL n=15, Spleen n=13). **C)** Frequency of QFL CD8SP T cells out of total CD8SP T cells in the indicated compartments of wild type mice. For tetramer enriched samples, frequencies out of CD8 T cells were determined by back-calculating to the unenriched samples (Thymus n=14, IEL n=15, Spleen n=13. Greyed area represents the range of frequencies observed for naïve conventional CD8SP T cells^39^. **D)** Ratio of CD5 gMFI of QFL CD8SP T cells from QFLTg mice (n=9) (Green) (Gated: QFL tetramer^+^CD24^-^ CD8α^+^CD4^-^) and non-transgenic mice (n=17) (Blue) (tetramer enriched and gated: QFL tetramer^+^ Vα3.2^+^ CD8α^+^CD4^-^) over conventional CD8SP CD5 gMFI. **E)** Representative plots of CD8β and CD8α expression on mature QFL CD8SP thymocytes from QFL TCR transgenic mice (QFLTg) (n=15) (Gated: QFL tetramer^+^CD24^-^CD8α^+^CD4^-^, as in Supplementary Fig. 1), non-transgenic tetramer enriched (n=32) (Gated: QFL tetramer^+^ and Vα3.2^+^ CD8α^+^CD4^-^) thymi. Conventional CD8SP (Gated: CD8α^+^CD4^-^) from unenriched non-transgenic thymi are shown for comparison. Dot plots show compiled data from QFLTg (green) and non-transgenic QFL tetramer^+^ thymocytes (blue). The average value for conventional CD8SP is indicated by dashed line. **F)** Representative histogram of CD44 expression on QFL CD8SP T cells from splenocytes of QFLTg mice (n=7) (Green), and non-transgenic mice (n=32) (Blue) (QFL tetramer enriched and gated: TCRβ^+^B220^-^QFL tetramer^+^Vα3.2^+^CD8α^+^CD4^-^), and conventional CD8SP T cells (Grey histogram). Dot plots show compiled data for QFL T cells from transgenic and non-transgenic spleen. The average value for conventional CD8+ splenocytes is indicated by dashed line. Error bars= Standard error of mean. Statistical analyses: One way ANOVA, followed by Tukey’s multiple comparison test. # P value <0.05, ## P value <0.005, ### P value <0.0005, #### P value <0.00001. One way ANOVA comparing samples to conventional CD8 T cells (shown below sample name). * P value <0.05, ** P value <0.005 *** P value<0.0005. Comparisons that are not statistically significant are not marked by a symbol.

As a complimentary method to characterize QFL T cells, we developed a TCR transgenic mouse that expresses the semi-invariant QFL TCR α-and β-chain (Vα3.2Jα21, Vβ1Dβ1Jβ2-7) used by a predominant clone ^37^, henceforth referred to as QFLTg. As expected, mature (CD24-) thymocytes expressing the QFL TCR were predominantly CD8 single positive (SP) (Supplementary Fig. 1A). DP thymocytes from transgenic mice expressed relatively low levels of the QFL TCR, while CD4SP thymocytes expressed moderately high levels of the TCR but remained mostly immature (CD24+) (Supplementary Fig. 2A). Expression of the QFL TCR transgene largely repressed endogenous TCRα expression, as indicated by the low levels of endogenous Vα2 on mature CD8SP T cells in both thymus and spleen (Supplementary Fig. 2B-C). In contrast, the frequency of Vα2 usage in CD4 SP thymocytes and splenocytes from QFL transgenic mice was similar to non-transgenic mice, confirming that they underwent positive selection using endogenous TCR rather than the QFL TCR. We also used thymocytes and splenocytes from QFLTg mice to confirm the specificity of the QFL TCR for the FL9-Qa1^b^ complex. As expected, the majority of QFL tetramer+ thymocytes and splenocytes did not stain with tetramers in which Qa1^b^ was loaded with the predominant QDM peptide (Supplementary Figure 2D). The small population of QFL tetramer+ cells that did stain with the QDM tetramer also expressed the QDM/Qa1^b^ receptor NKG2A, an NK receptor that is also expressed by activated T cells. The co-staining of QDM tetramer and NKG2A suggests TCR independent binding of QDM tetramer on QFL T cells and serves as a positive control for QDM tetramer staining. Taken together this data validates the efficacy of our TCR transgenic system and confirms the specificity of the QFL TCR for the FL9/Qa1^b^ complex.

Previous studies showed that QFL T cells respond to a self-peptide presented by Qa1^b27^. In addition, splenic QFL T cells from wild type mice display an antigen experienced phenotype, suggesting that they may receive strong TCR signals during their development in the thymus. To test this notion, we examined expression of CD5, a marker which positively correlates with self-reactivity ^41–44^. As predicted, we observed that CD5 is elevated in QFL CD8SP thymocytes from QFLTg and non-transgenic mice compared to conventional CD8SP T cells (Fig. 1D). In addition, QFL CD8SP thymocytes from non-transgenic mice showed slight but detectable downregulation of CD8β compared to conventional CD8SP T cells, whereas QFL CD8SP thymocytes from QFLTg mice showed more pronounced CD8β downregulation (Fig. 1E). This modulation of CD8β expression has been associated with thymocyte self-reactivity and agonist selection^45,46^. QFL CD8SP thymocytes from QFLTg mice also showed elevated levels of several markers associated with agonist selection, such as the transcription factors PLZF ^47,48^ and Tbet (Supplementary Fig. 3). On the other hand, splenic, but not thymic, QFL CD8SP T cells express elevated levels of CD44, a marker associated with antigen experience (Fig. 1F) (Supplementary Fig. 3A, C). In addition, QFL CD8SP thymocytes from QFLTg mice did not show detectable upregulation of PD1 or α4β7, markers that are expressed by a subset of thymic IEL precursors (Supplementary Fig. 3A, C) ^21^. Taken together these data suggest that QFL T cells experience relatively strong TCR stimulation and undergo agonist selection during their development in the thymus.

### QFL T cell development in absence of Qa1^b^ or classical MHC I

Some agonist selected T cell populations, such as regulatory T cells require a separate positive selection interaction prior to undergoing agonist selection^6^. Previous reports that QFL T cells are detectable in the spleen of mice lacking Qa1^b^, but undetectable in mice lacking β2m^27^, a subunit of MHC I which is required for proper folding and surface expression of both classical MHC Ia and Qa1 ^49^, suggested the possibility that QFL T cells require positive selection on classical MHC Ia. To test this hypothesis, we generated K^b^D^b^KO mice and compared the number of QFL T cells in the thymus and spleen to that of WT and Qa1^b^KO mice (Fig. 2A, Supplemental Fig. 4A-B). The QFL CD8SP thymocytes were slightly reduced in K^b^D^b^KO and Qa1^b^KO relative to wild type mice but were undetectable in β2mKO mice (Fig. 2A). Similar results were obtained with QFLTg mice, with substantial numbers QFL thymocytes and splenocytes found in the absence of Qa1^b^ or K^b^D^b^, but not in the absence of β2m (Fig. 2B). Importantly, the QFL CD8SP thymocytes and splenocytes from QFLTg, QFLTgQa1^b^KO mice have negligible expression of endogenous Vα2 (Supplemental Fig. 2B-C), confirming that these cells were positively selected using the QFL TCR. Classical MHC Ia D^b^ is the source for the QDM peptide that is the predominant peptide bound to Qa1^b^ in ERAAP sufficient cells, raising the possibility that loss of D^b^ could indirectly impact QFL T cell development by altering the peptide displayed on Qa1^b^. However, QFL CD8SP thymocytes and splenocytes were found in similar numbers in K^b^KO, D^b^KO, and K^b^D^b^KO mice, arguing against this possibility (Supplementary Fig. 4C). Altogether, these data suggest that neither classical MHC Ia, nor Qa1^b^, are required for QFL T cell positive selection, although both may contribute to the efficiency of the process. Interestingly, CD8SP T cells in spleens of K^b^D^b^KO mice exhibit a higher frequency of Vα3.2^+^ cells compared to WT or Qa1^b^KO mice (Supplementary Fig. 4D), indicating that this V segment is preferentially used by T cells reactive to MHC Ib molecules.

**Figure 2:**
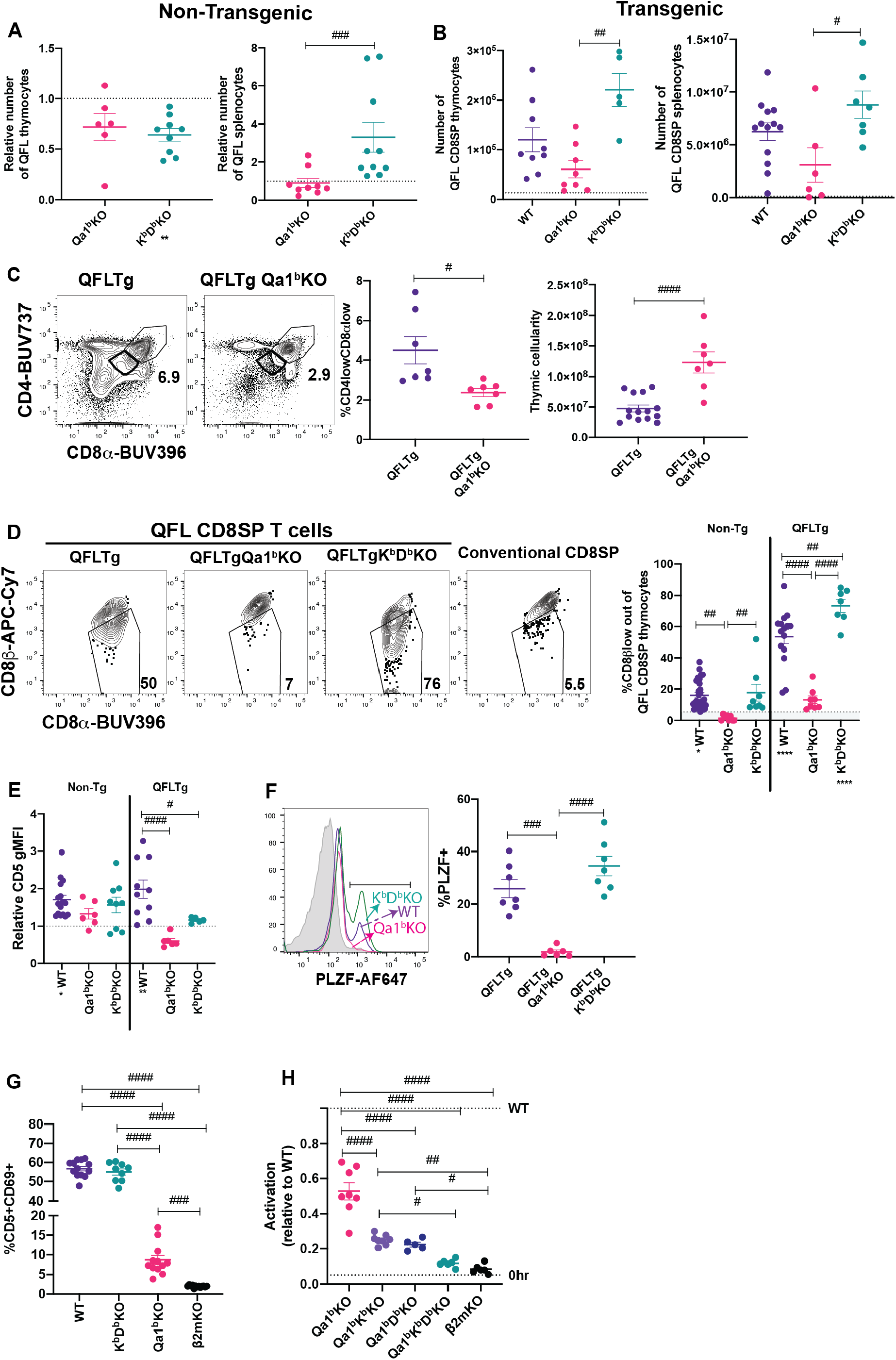
MHC requirements for QFL T cell development. **A)** Relative number of QFL T cells in thymi (tetramer enriched and gated: QFL tetramer^+^Vα3.2^+^CD8α^+^CD4^-^) or spleen (tetramer enriched and gated: TCRβ^+^QFL tetramer^+^Vα3.2^+^ CD8α^+^CD4^-^) of non-transgenic mice of the indicated genotype. To correct for variation in the efficiency of tetramer enrichment, data are normalized to the number of QFL T cells recovered from a wild type mouse analyzed in the same experiment (represented by dotted line) (Thymus Qa1^b^KO n=6, K^b^D^b^KO n=9) (Spleen Qa1^b^KO n=9, K^b^D^b^KO n=10. **B)** Number of QFL CD8SP T cells in the thymi (Gated: QFL tetramer^+^ CD24^-^CD8α^+^CD4^-^)(WT n=9, Qa1^b^KO n=8, K^b^D^b^KO n=5) or spleens (Gated: TCRβ^+^QFL tetramer^+^Vα3.2^+^ CD8α^+^CD4^-^) (WT n=13, Qa1^b^KO n=6, K^b^D^b^KO n=7) of QFLTg mice crossed to the indicated gene knock out strains. Dotted line represents the limit of detection which was based on β2mKO control. **C)** Representative plots of CD4 and CD8α expression on QFL thymocytes from QFLTg and QFLTg Qa1^b^KO mice. Dot plot shows compiled data for % of CD4lowCD8αlow in individual mice (Gated: live cells). Panel to the right shows thymus cellularity from QFLTg and QFLTg Qa1^b^KO mice. Each dot represents an individual mouse (n=8 for all conditions). **D)** Representative plots of CD8β and CD8α expression on QFL CD8SP thymocytes (Gated: QFL tetramer^+^CD24^-^ CD8α^+^CD4^-^) from QFLTg, QFLTg Qa1^b^KO and QFLTg K^b^D^b^KO mice. Conventional CD8+ thymocytes from wild type mice (Gated:CD8α^+^CD4^-^) are shown for comparison. Graph shows % of CD8β low out of QFL CD8SP T cells in thymi of non-transgenic (WT n=26, Qa1^b^KO n=8 and K^b^D^b^KO n=8) or QFLTg (WT n=15, Qa1^b^KO n=8 and K^b^D^b^KO n=7) mice of the indicated genotypes. Dotted line indicated the average value (5.2%) for conventional CD8SP (Gated: CD8α^+^CD4^-^) from wild type mice. **E)** CD5 expression on QFL CD8SP thymocytes from either non transgenic (WT n=17, Qa1^b^KO n=6 and K^b^D^b^KO n=9) or QFLTg (WT n=10, Qa1^b^KO n=6 and K^b^D^b^KO n=5) mice of the indicated genotypes. Graph shows gMFI of CD5 expression of QFL T cells normalized to the gMFI of conventional CD8SP thymocytes from wild type mice analyzed in the same experiment. **F)** Representative histogram of PLZF expression in QFL CD8SP T cells from QFLTg mice of the indicated genotypes (QFLTg n=7, QFLTg Qa1^b^KO n=6 and QFLTgK^b^D^b^KO n=7). Grey histograms represent conventional CD8SP (Gated:CD8α^+^CD4^-^). **G)** Pre-selection QFL thymocytes (preQFLTg) (from QFLTg β2mKO mice) were co-cultured with Bone Marrow Derived Dendritic cells (BMDC) from the indicated mouse strains. Compiled data of preQFLTg DP thymocyte (Gated: QFL tetramer^+^Vα3.2^+^CD4^+^CD8α^+^) expression of CD5 and CD69 after 24 hours of co-culture. Dot plots show compiled data from three experiments, with each dot representing a sample from an individual culture well (WT n=13, Qa1^b^KO n=12, K^b^D^b^KO n=9 and β2mKO n=13). **H)** preQFLTg thymocytes were co-cultured for 24Hrs with either parental (WT) DC2.4 cells or DC2.4 cells in which genes encoding the indicated MHC molecules were knocked out using CRISPR/Cas9 editing. The percentage of CD5^+^CD69^+^preQFLTg (Gated: QFL tetramer^+^Vα3.2^+^) thymocytes from each co-culture condition (Qa1^b^KO, Qa1^b^K^b^KO, Qa1^b^D^b^KO, Qa1^b^K^b^D^b^KO, and β2mKO) was normalized to the percentage of CD5^+^CD69^+^preQFLTg thymocytes co-cultured with WT DC2.4 cells, where WT=1 (Top dotted line). 0Hrs (Bottom dotted line) negative control represents the percentage of CD5^+^CD69^+^preQFLTg thymocytes before addition to co-culture. Dot plots show compiled data from three experiments, with each dot representing a sample from an individual culture well (Qa1^b^KO n=8, Qa1^b^K^b^KO n=8, Qa1^b^D^b^KO n=5, Qa1^b^K^b^D^b^KO n=5, and β2mKO n=6). Error bars= Standard error of mean. Statistical analysis: Kruskal-Wallis test followed by Dunn’s test multiple comparison test. For comparisons across displayed samples (shown above dots) # P value, <0.0332, ## P value <0.0021, ### P value<0.0002, #### P value<0.00001. For comparisons to wild type samples (panel A), or to conventional CD8SP (panel B) (shown below sample name) * P value, <0.0332, ** P value <0.0021, *** P value<0.0002, **** P value<0.00001. One way ANOVA followed by Tukey’s multiple comparison test. For comparisons across displayed samples (shown above dots) # P value, <0.05, ## P value <0.005, ### P value<0.0005, #### P value<0.00001. For comparisons to conventional CD8SP (panel D and E) (shown below sample name) * P value <0.05, ** P value <0.005 *** P value<0.0005. Comparisons that are not statistically significant are not marked by a symbol.

Because Qa1^b^ presents agonist FL9 peptide to QFL T cells, we hypothesized that expression of Qa1^b^ might lead to agonist and negative selection of QFL thymocytes. In support of this, DP thymocytes in QFLTg (Qa1^b^ sufficient) mice exhibit reduced cellularity and a “DP^lo^” phenotype associated with strong TCR signals ^19,50–52^ whereas QFLTg Qa1^b^KO mice express normal levels of CD4 and CD8α (Fig. 2C). In addition, QFL CD8SP thymocytes from Qa1^b^ sufficient, but not Qa1^b^KO mice, displayed CD8β downregulation, PLZF expression, and elevated CD5 expression compared to conventional mature CD8SP thymocytes (Fig. 2D-F). In contrast, in the absence of classical MHC I (K^b^D^b^KO mice) QFL CD8SP T cells showed strong downregulation of CD8β expression, maintained PLZF expression and showed a slight reduction in CD5 expression compared to WT mice (Fig. 2D-F). In the periphery, QFL T cells lost their antigen experienced phenotype in absence of Qa1^b^, but not in absence of classical MHC I (Supplemental Fig. 4E). These data suggest that Qa1^b^ is required for agonist selection of QFL T cells, although positive selection of QFL T cells can be driven by an alternative MHC 1 molecule.

### QFL T cells recognize an alternative ligand on Qa1^b^KO APCs

To further explore the ligand-specificity of the QFL TCR we took advantage of the observation that MHC-naïve DP thymocytes are highly sensitive to in vitro TCR stimulation ^53,54^. We examined expression of activation markers on pre-selection QFLTg (preQFLTg) thymocytes from a β2mKO background after co-culture with bone marrow derived dendritic cells (BMDC) isolated from mice lacking either Qa1^b^, K^b^D^b^ or β2m. PreQFLTg thymocytes showed stronger upregulation of the activation markers CD69 and CD5 upon 24-hour co-culture with WT and K^b^D^b^KO, compared to Qa1^b^KO, BMDC (Fig. 2G) (Supplementary Fig. 5A). This is consistent with the hypothesis that Qa1^b^ is the primary MHC ligand for the QFL TCR. Interestingly, preQFLTg thymocytes showed a modest activation when co-cultured with Qa1^b^KO BMDCs; this activation was significantly more compared to co-culture with β2mKO BMDCs (Fig. 2G). A similar pattern of reactivity was observed when preQFLTg were cultured in thymic slices derived from WT, Qa1^b^KO and β2mKO mice (Supplementary Fig. 5B-C). This is consistent with the development of QFL T cells in Qa1^b^KO mice, and suggests that the QFL TCR is cross-reactive with an alternative β2m-utilizing molecule, most likely classical MHC I.

To specifically test whether the QFL TCR cross-reacts with classical MHC 1 (K^b^D^b^), we used the DC-like cell line DC2.4 as a stimulator cell for preQFLTg thymocytes (Fig. 2H). The response of preQFLTg thymocytes to DC2.4 cells was abolished by CRISPR/Cas9 mediated gene knock out of β2m and partially reduced by loss of Qa1^b^, paralleling the results from stimulation with BMDC, and pointing to the recognition of an alternative MHC-1 ligand in this system (Fig. 2G-H). Interestingly, triple KO of Qa1^b^, K^b^, and D^b^ in DC2.4 cells reduced activation close to background levels, whereas double knock out of Qa1^b^ with either K^b^, or D^b^ led to stimulation that was intermediate between the triple KO and Qa1^b^ KO cell lines. Together these data suggest that the QFL TCR has low level reactivity with both K^b^ and D^b^ and suggests that these two classical MHC-1 molecules may contribute to the positive selection of QFL T cells in Qa1^b^KO mice.

### QFL T cell selection by hematopoietic and non-hematopoietic cells

While conventional αβT cells undergo positive selection by recognition of MHC molecules on thymic epithelial cells, MAIT cells and iNKT cells undergo selection via interactions with hematopoietic cells ^4,5^. To investigate the cell type requirements for selection of QFL T cells, we generated reciprocal bone marrow chimeric mice in which either the donor cells or the host cells are β2mKO, and therefore lack surface expression of Qa1, as well as the classical MHC I molecules H2-D, H2-K (Supplementary Fig. 6). Interestingly, comparable numbers of QFL T cells were found in the thymus and spleen of the β2mKO>WT and WT>β2mKO chimeric mice (Supplementary Fig. 6B-C), implying that QFL T cell development could occur efficiently on either non-hematopoietic or hematopoietic cells. To confirm these results, we generated reciprocal β2mKO chimeras using donor cells that expressed the QFL TCR transgene (Fig. 3A). While there was some reduction in QFL T cell number in the thymus of β2mKO>WT compared to WT>β2mKO and wild type control chimeras (Fig. 3B), similar numbers of QFL T cells were found in the spleen (Fig. 3C). Thus, QFL T cell development is not strictly dependent on either non-hematopoietic or hematopoietic expression of MHC I.

**Figure 3:**
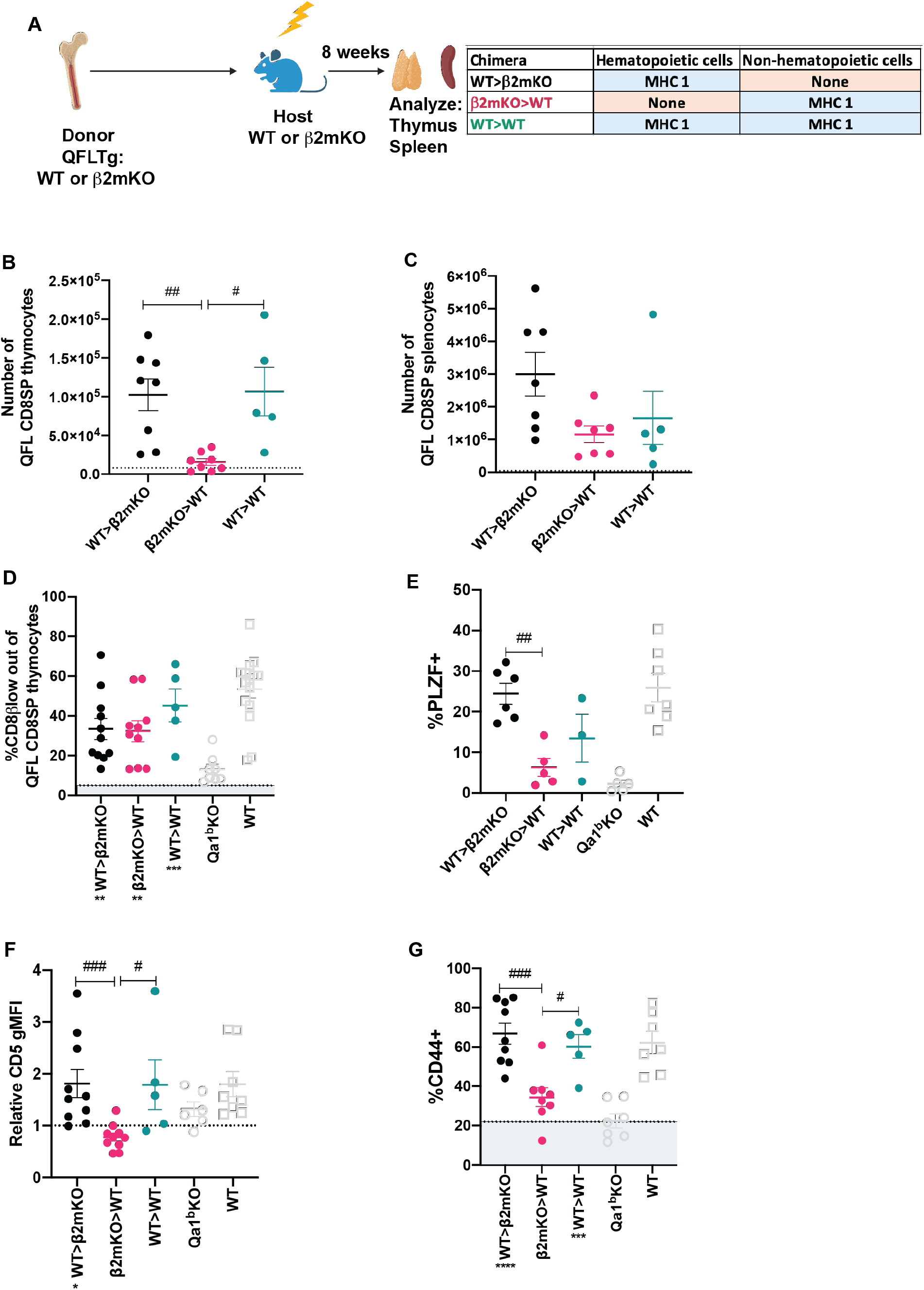
Requirement for hematopoietic versus non-hematopoietic cell MHC I expression for QFL thymic selection. **A)** Diagram of experimental design. QFLTg or QFLTg β2mKO mice were used as bone marrow donors to reconstitute irradiated β2mKO or wild type hosts in order to restrict MHC I expression to hematopoietic or non-hematopoietic cells. **B-C)** Absolute numbers of QFL CD8SP T cells in the indicated chimeric mice in **B)** Thymus (gated: QFL tetramer^+^CD24^-^CD8α^+^CD4^-^) (WT>β2mKO n=8, β2mKO>WT n=8, WT>WT n=5) and **C)** Spleens (gated: TCRβ^+^QFL tetramer^+^Vα3.2^+^CD8α^+^CD4^-^)(WT>β2mKO n=7, β2mKO>WT n=7, WT>WT n=5). Dotted line represents the limit of detection which was based on β2mKO>β2mKO control. **D)** Downregulation of CD8β on QFL CD8SP thymocytes of the indicated chimeric mice (WT>β2mKO n=11, β2mKO>WT n=10, WT>WT n=6). Dotted line represents the average for conventional CD8SP (Gated: CD8α^+^CD4-) from unenriched non-transgenic thymi (5.2%). **E)** Quantification of PLZF expression in QFL CD8SP thymocytes from the indicated chimeric (WT>β2mKO n=6, β2mKO>WT n=5, WT>WT n=3). **F)** CD5 expression on QFL CD8SP thymocytes of the indicated chimeric mice. Graph shows gMFI of CD5 expression of QFL thymocytes normalized to the gMFI of conventional CD8SP thymocytes from wild type mice analyzed in the same experiment (WT>β2mKO n=10, β2mKO>WT n=10, WT>WT n=5). **G)** Quantification of CD44 expression of QFL CD8SP T cells (Gated: TCRβ^+^QFL tetramer^+^Vα3.2^+^CD8α^+^CD4^-^) from: WT>β2mKO (Black dots), β2mKO>WT (Magenta dots), WT>WT (Teal dots) chimeric spleens and Qa1^b^KO (Light blue dots) spleens. Dotted line represents the average value for conventional CD8SP (Gated: TCRβ^+^CD8α^+^CD4^-^) (22%) (WT>β2mKO n=9, β2mKO>WT n=8, WT>WT n=5). In D-G, data from intact QFLTg and QFLTg Qa1^b^KO mice from Figure 2D is included for comparison. Error bars= Standard error of mean. Statistical analysis: Kruskal-Wallis test followed by Dunn’s test multiple comparison test (Panel B and C). For comparisons across displayed samples (shown above dots) # P value, <0.0332, ## P value <0.0021, ### P value<0.0002, #### P value<0.00001. One way ANOVA followed by Tukey’s multiple comparison test. # P value, <0.05, ## P value <0.005, ### P value<0.0005, #### P value<0.00001. One way ANOVA comparing samples to conventional CD8 T cells (shown below sample name). * P value <0.05, ** P value <0.005 *** P value<0.0005. Comparisons that are not statistically significant are not marked by a symbol.

We also examined whether QFL T cells that are selected exclusively by hematopoietic or non-hematopoietic cells retained their agonist selected phenotype. QFL thymocytes exhibited comparable CD8β downregulation but decreased PLZF expression when MHC I was restricted to non-hematopoietic cells (Fig. 3D-E, Supplementary Fig. 6D). Similarly, expression of CD5 was decreased when MHC I was restricted to non-hematopoietic cells (Fig. 3F, Supplementary Fig. 6E). In the periphery, QFL T cells in chimeric mice that lacked MHC I on hematopoietic cells did not display elevated expression of CD44 (Fig. 3G, Supplementary Fig. 6F). Overall, the thymic phenotype of QFL T cells in β2mKO>WT chimeras is similar, but not identical, to that observed in Qa1^b^KO mice (Fig. 2D-F). These data suggest that Qa1^b^ on both hematopoietic and non-hematopoietic cells contribute to agonist selection, with hematopoietic cells playing the predominant role.

### Impact of agonist selection on QFL T cell function

To test how agonist selection impacts the functionality of QFL T cells, we compared QFL T cells that arose in the presence or absence of Qa1^b^ for their ability to respond *in-vitro* to ERAAPKO splenocytes. QFL T cells from QFLTg showed extensive upregulation of activation markers and increased proliferation in response to stimulation compared to QFL T cells from QFLTg Qa1^b^KO mice (Fig. 4A-B). This implies that agonist selection on Qa1^b^ led to greater functional responsiveness, which could reflect greater functionality on a per cell basis, or an increased frequency of functional cells within the population. Since agonist selection partially correlates with selection on hematopoietic cells (Fig. 3, Supplementary Fig. 6), we also examined QFL splenocytes from reciprocal β2mKO and wild type bone marrow chimeric mice as a further test of the impact of agonist selection on function. QFL T cells from QFLTg>β2mKO mice (Hematopoietic cell (HC) selected) responded more robustly to ERAAPKO APCs compared to cells from QFLTg β2mKO>WT mice (non-HC selected) (Fig. 4A-B). Thus, QFL T cells that develop in the absence of Qa1^b^, or in the absence of hematopoietically expressed MHC I, exhibit reduced functionality.

**Figure 4:**
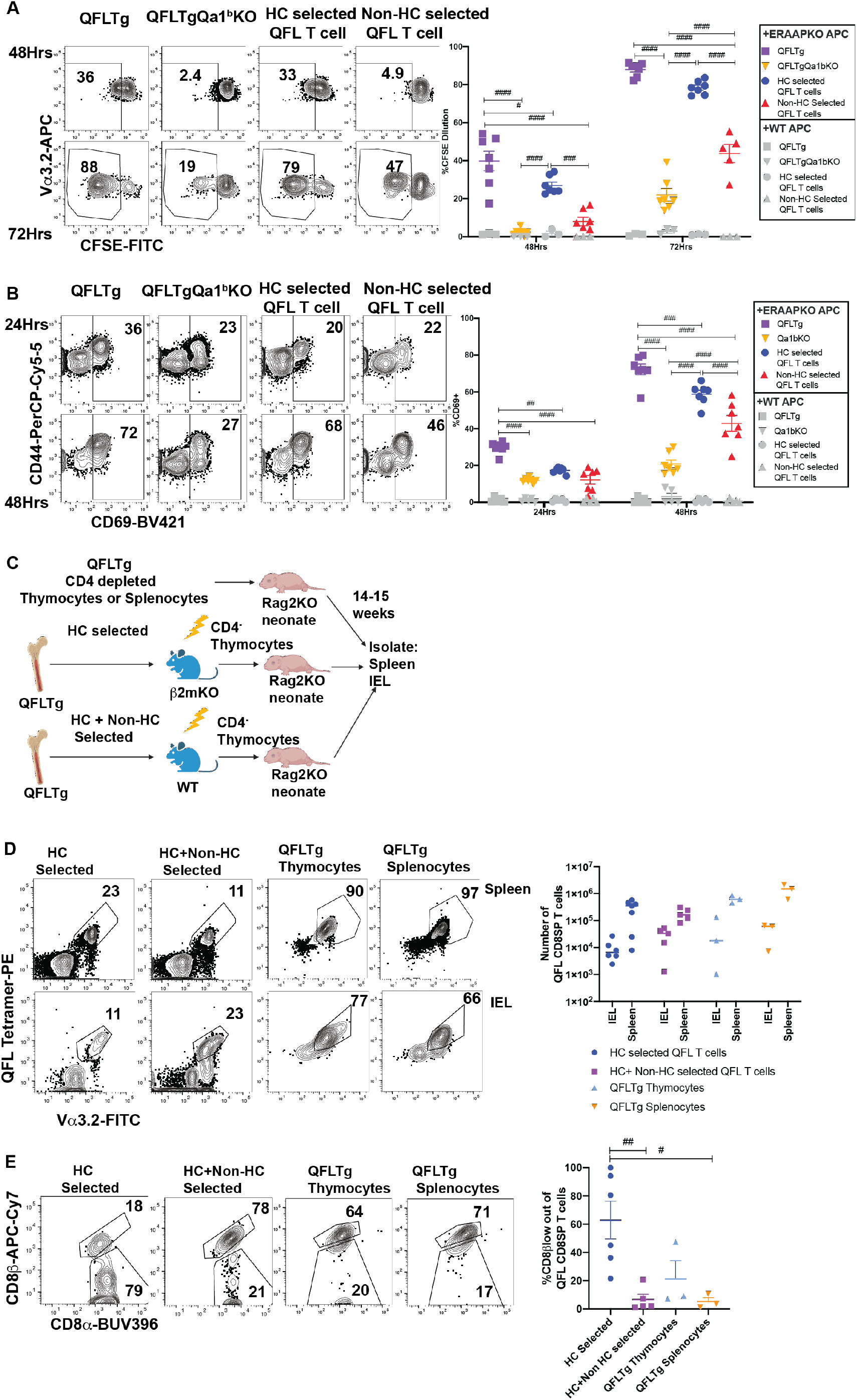
Agonist selected QFL T cells respond rapidly to antigen exposure and home to the IEL compartment. **A-B)** QFL CD8SP T cells from QFLTg or QFLTgQa1^b^KO mice or from QFLTg>β2mKO (Hematopoietic cell (HC) Selected) or QFLTgβ2mKO>WT (non-HC selected) bone marrow chimeric mice were labeled with CFSE and co-cultured with splenocytes from ERAAP KO mice, and analyzed after 24, 48, or 72 hours of co-culture. Representative plots and quantification of CFSE dilution **(A)** or CD69 surface expression **(B)** on QFL CD8SP T cells (Gated: TCRβ^+^B220^-^QFL tetramer^+^ Vα3.2^+^CD8α^+^). Compiled data of two experiment (n=7 for all conditions). **C)** Experimental design: Bone marrow chimeric mice were generated using QFLTg bone marrow donors and hosts deficient or sufficient for β2m, and CD4- depleted thymocytes from the chimeric mice were injected into Rag2KO neonates (Figure 3A). For comparison, CD4-depleted QFLTg thymocytes or splenocytes were transferred into Rag2KO neonates. The spleen and small intestinal IEL compartments were analyzed 14-15 weeks post injection. **D)** Representative flow cytometry plots of QFL tetramer and Vα3.2 from spleen or IEL of the indicated transferred Rag2KO mice (Gated: Donor^+^TCRβ^+^) (HC selected n=7, HC+nonHC selected n=5, QFLTg thymocytes n=3, and QFLTg splenocytes n=3). Compiled data show the number of QFL tetramer^+^Vα3.2^+^CD8α^+^ cells recovered from each sample. Each dot represents an individual mouse **E)** Representative plots of surface expression of CD8α and CD8β on QFL CD8SP T cells (Gated: TCRβ^+^QFL tetramer^+^Vα3.2^+^CD8α^+^) isolated from the IEL compartment of the indicated transferred Rag2KO mice (HC selected n=6, HC+nonHC selected n=5, QFLTg thymocytes n=3, and QFLTg splenocytes n=3). Compiled data shows %CD8β low out of QFL CD8SP IEL T cells recovered from indicated transferred Rag2KO mice. Each dot represents an individual mouse. Error bars= Standard error of mean. Statistical analyses: Two-way ANOVA followed by Tukey’s multiple comparison test comparing samples to each other in their respective time point. # P value, <0.05, ## P value <0.005, ### P value<0.0005, #### P value<0.00001. student’s T test. * P value, <0.05, **P value <0.005, *** P value<0.0005, ****P value<0.00001. Comparisons that are not statistically significant are not marked by a symbol.

### QFL thymocytes and splenocytes can populate the intestinal epithelial compartment

QFL thymocytes display an agonist selected phenotype that is enhanced by hematopoietic expression of MHC I. Given that agonist selection in the thymus can give rise to natural intraepithelial lymphocytes, we considered that QFL thymocytes might represent a population of IEL precursors. To test this idea, we injected Rag2KO neonatal mice with CD4-depleted QFL thymocytes from HC-only selected (QFLTg>β2mKO) or both HC and non-HC selected (QFLTg>WT) chimeric mice (Fig. 4C). For comparison, we also injected Rag2KO neonatal mice with CD4-depleted thymocytes or splenocytes from intact QFLTg mice. QFL T cells were found in similar numbers in the spleen and IEL compartment of the SI for all 4 donor populations (Fig. 4D). Interestingly, IEL T cells derived from HC-only selected QFL thymocytes showed more pronounced downregulation of CD8β (∼50%) when compared to IEL T cells derived from HC+ non-HC selected QFL T cells (∼10%) (Fig. 4E). Altogether, these data indicate that QFL thymocytes and splenocytes contain IEL precursors, and that selection by hematopoietic cells favors CD8β downregulation in QFL IELs.

## Discussion

Most studies of unconventional T cells have focused on 2 prominent populations, MAIT (MR1-restricted) and iNKT (CD1d restricted) cells, and much less is known about the development of T cells restricted to other MHC Ib molecules. Moreover, while it is known that non classical MHC molecules contribute substantially to the CD8αα natural IEL compartment^13,15,53^, and there is evidence that thymic mature CD4-CD8- (DN) cells contain IEL precursors ^20,21,55,56^, it is unclear whether all natural IEL develop via a mature DN pathway. Here we have used both QFL TCR transgenic mice and FL9-Qa1^b^ tetramer staining of non-transgenic mice to investigate the development of a population of self-reactive Qa1^b^ restricted cells known as QFL T cells. QFL T cells are found in circulation as both naïve and memory phenotype CD8αβ T cells, and in the IEL compartment as both CD8αα and CD8αβ cells, whereas mature QFL thymocytes are predominantly CD8αβ+CD4- and show signs of agonist selection. QFL T cells have a more relaxed requirement for positive selection compared to conventional CD8 T cells, requiring β2m on either hematopoietic or non-hematopoietic cells, but neither the restricting molecule Qa1^b^, nor MHC Ia for positive selection. However, QFL thymocytes do require Qa1^b^ for agonist selection and full functionality. Our data highlight the promiscuous requirements for positive selection of a Qa1 restricted T cell population and identify an alternative CD8αβ+CD4- pathway for development of CD8αα IELs.

The flexible thymic development of QFL T cells parallels their ability to give rise to T cells with both conventional and unconventional properties. Unconventional MAIT and iNKT cells require selection by their restricting MHC molecules on hematopoietic cells, giving rise to T cells that migrate directly to tissues and exhibit preformed effector program. On the other hand, conventional CD8 T cells require selection by their restricting MHC Ia molecules on thymic epithelial cells, producing circulating naïve T cells that lack effector programing. QFL T cells appear to have the option to develop by either of these pathways, with selection by Qa1^b^ on hematopoietic cells leading to a more unconventional phenotype, and selection via an alternative MHC I ligand giving rise to T cells that resemble conventional CD8 T cells.

Interestingly, the ability to be selected on either hematopoietic or non-hematopoietic cells in the thymus has been reported both for another Qa1 restricted T cell population ^9^, as well as a T cell population restricted to the MHC Ib molecule H2-M3 ^8^. While the M3 restricted cells required M3 expression for thymic selection, hematopoietic selection led to T cells with more unconventional functional properties compared to non-hematopoietic selection. Thus, a flexible pattern of thymic selection leading to alternative functional programs may be a general feature of T cell reactive to Qa1 and H2-M3.

The IEL compartment harbors 2 distinct types of αβTCR+CD8+ T cells: “induced” CD8αβ T cells that are derived from conventional CD8 T cells following encounter with foreign antigen and differentiation into tissue resident memory T cells, and “natural” CD8αα IEL that are directed into an IEL program in the thymus due to their high self-reactivity^16,17^. The observation that the same TCR clone can give rise to both types of IEL blurs the distinction between these two types of cells. Previous studies of natural IEL development have suggested a pathway in which some DP thymocytes that receive strong TCR signals escape clonal deletion by downregulating CD4 and CD8 to give rise to mature CD4-CD8- IEL precursors (IELp), that can eventually migrate to the gut and upregulate CD8αα ^17,21^. Our data are consistent with an alternative pathway for IEL development in which agonist selection leads to a mature CD8αβ+CD4- thymic IELp. While the ability of CD4-depleted thymocytes to repopulate the IEL compartment of Rag2KO mice does not rule out a contribution from CD4-CD8α- precursors, the absence of detectable CD4-CD8α- QFL thymocytes in non-transgenic mice, as well as the agonist phenotype of thymic QFL CD8SP, strongly suggests that the CD4-CD8αβ cells are the relevant thymic precursor population. Moreover, the partial downregulation of CD8β observed on mature QFL thymocytes and on some QFL IEL T cells, suggests that CD8β expression may be unstable in QFL cells, leading to partial or full downregulation once they arrive in the IEL compartment. The suggestion is also consistent with the observation that splenic QFL T cells, which are uniformly CD4-CD8αβ, can give rise to IELs expressing intermediate levels of CD8β upon transfer into Rag2KO recipients. Altogether, these data are in line with earlier studies of thymocytes agonist selection in organ culture that also implicated mature CD8SP as a thymic precursor to natural IEL^45,46^. A detailed understanding of the developmental pathways and signals involved in QFL IEL T cell development awaits further investigation.

Our data, together with published observations, support the notion that MHC-E restricted CD8 T cells are generally cross-reactive. Using an *in vitro* stimulation assay with pre-selection QFL thymocytes, which reads out relatively weak TCR signals compatible with positive selection ^53,54^, we found that QFL thymocytes can respond to Qa1^b^KO APCs, but not to APC lacking Qa1^b^ as well as both K^b^ and D^b^ classical MHC1a molecules. Thus, cross-reactivity to classical MHC-1 molecules likely accounts for the positive selection of QFL thymocytes in the absence of Qa1^b^. In addition, another Qa1^b^ restricted clone was shown to cross react with an MHC Ia molecule ^57^, although it was dependent on Qa1^b^ for its positive selection ^9^. The MHC-E restricted response to a CMV-vectored HIV vaccine showed extremely broad reactivity, with detectable responses to 4 epitopes for every 100 amino acids ^58^. In this regard, it is intriguing that QFL T cells show strong preferential use of Vα3.2 (encoded by TRAV9N/D-3) ^37^. Vα3.2 is preferentially used by CD8 compared to CD4 T cells ^59^ and has been suggested to be inherently reactive to MHC I ^60^. In addition, Vα3.2 is enriched in a subset of natural IELs ^21^, and is used by another Qa1^b^-restricted CD8 T cell clone ^9^. Moreover, we found that the frequency of Vα3.2+ CD8 T cells is substantially increased in K^b^D^b^KO mice (Supplementary Figure 4D). Altogether, these observations suggest that Vα3.2 may work together with Qa1^b^, and perhaps other non-classical MHC I molecules, to generate self-reactive T cells with a propensity to give rise to memory phenotype and natural IEL T cells.

If MHC-E reactive CD8 T cells are inherently cross-reactive, how do they escape negative selection in the thymus? While thymocyte intrinsic mechanisms, such as downregulation of CD4 and CD8 may contribute ^52^, it is interesting to consider how properties of the MHC molecules may also play a role. In particular, MHC-E molecules tend to be expressed at lower levels on the cell surface compared to MHC Ia molecules ^9,61,62^, a property that may be linked to their atypical peptide presentation pathway ^63,64^ and/or low surface stability ^61^. In addition, MHC-E molecules predominantly express a single self-peptide derived from MHC Ia leader peptides ^65,66^, and may not present a large array of self-peptides in healthy cells. Indeed, it has been proposed that MHC-E molecules may monitor alterations in the MHC Ia peptide presentation pathway that occur upon viral infection or cellular transformation^10,27,29^, changes which may be mimicked by conditions of cellular stress. According to this notion, MHC-E restricted T cells may undergo rare or transient encounters with high affinity self-peptide-MHC-E complexes on stressed cells during their development in the thymus, allowing them to experience agonist selection signals while avoiding negative selection.

## Acknowledgements

We thank Kartoosh Heydari, H. Nolla, and A. Valero of the UC Berkeley Cancer Research Lab for help with flow cytometry. We thank members of the Shastri, Coscoy, Raulet, and Robey lab for helpful discussions. Funding was provided by the National Institutes of Health (RO1AI149341). M.M. was supported by a National Science Foundation Graduate Research Fellowship and a diversity supplement to R01AI149341.

## Methods

### Mice

B6 (C57BL/6), B6 Ly5.1 (B6.SJL-*Ptprca Pepcb*/BoyJ) and Rag2-/- (B6(Cg)-Rag2tm1.1Cgn/J) mice were from Jackson Labs. Β2M-/- (B6.129-B2mtm1Jae N12) mice were from Taconic. Qa1^b^KO mice ^67^ were obtained from the Shastri lab. TCR transgenic mice specific for FL9-Qa1^b^ (QFL) and H-2K1/H-2D1-/- (K^b^D^b^KO) mice were generated in our lab (described below). All mice were bred in the UC Berkeley animal facility and all procedures were approved by the Animal Care and Use Committee (ACUC) of the University of California.

### Generation of the QFLTg mouse

The TCR alpha and beta chain sequences from the QFL specific BEKo8z hybridoma^27,37^ were cloned and amplified from the genomic DNA of the BeKoz Hybridoma. The TRAV9N/D-3 TCR alpha chain was cloned with the forward primer (5’ AAAACCCGGGCCAAGGCTCAGCCATGCTCCTGG) with an added XmaI cutting site at 5’ end of the DNA sequence and a reverse primer for TRAJ21 (5’ AAAAGCGGCCGCATACAACATTGGACAAGGATCCAAGCTAAAGAGAACTC) with an added Not1 cutting site at the 5’ end of the DNA sequence. The TCR beta chain was cloned with the forward primer (5’ AAAACTCGAGCCCGTCTGGAGCCTGATTCCA) with and added Xho1 cutting site at the 5’ end of the DNA and a reverse primer for TRBJ2-7 (5’ AAAACCGCGGGGGACCCAGGAATTTGGGTGGA) with a SacII cutting site flanking the 5’ end of the DNA sequence. The cloned TCR alpha chain was cloned into pTα cassette vector by inserting it between the Xmal and Not1 sites, while the TCR beta chains were cloned into pTβ cassette vector in between the Xhol and SacII sites ^68^. The ampicillin resistance gene was removed from pTα and pTβ cassette by EarI enzyme digest. The QFL transgenic mice were generated on the B6 background in the Cancer Research Laboratory Gene Targeting Facility at UC Berkeley under standard procedures. The QFL mice were maintained on the B6 background and bred once with B6 5.1 mice to generate (QFLTgxB65.1/2) background mice for use in experiments. Founder mice were identified by flow cytometry and PCR genotyping of tail genomic DNA using primers mentioned above.

### Generation of K^b^D^b^KO mice

K^b^D^b^KO were generated by the Gene Targeting Facility at UC Berkeley using Cas9/CRISPR-mediated gene targeting. The H-2K1 gene was targeted using an sgRNA (5’ GTACATGGAAGTCGGCTACG 3’) that aligned with the sense strand and the H-2D1 gene was targeted using an sgRNA (5’ AGATGTACCGGGGCTCCTCG 3’) that aligned with the antisense strand. Wild-type C57BL/6J mice were originally obtained from the Jackson Laboratories. Zygotes were obtained from super ovulated C57BL/6J females for CRISPR/Cas9 targeting knockout experiment. In brief, CRISPR mix (i.e., Cas9 protein and sgRNAs) was introduced to zygotes by electroporation as previously described ^69^. The embryos were then transferred to 0.5dpc pseudo pregnant females (CD-1, Charles River Laboratories) with oviduct transfer. When the pups were born, tails samples were collected for DNA extraction and genotyping. The resulting founder mice were identified by flow cytometry. The H-2K1 gene had a 2bp deletion (5’ TGCCTGGGCTTTCTGTGTCTCCCGCTCCCAATACTCGGGCCCCTCCTGCTCCATCCACCGC GCCCGCGGCTCATATCTCGGATTCTCCGCGTCGCTGTCGAAGCGCACGAACTCCGTGTCGTCCACGT— CCGACTTCCATGTACCGGGGCTCCCCGAGGCCGGGCCGGGACACGGCGGTGACGAAATACCTCAA 3’) where the sgRNA targeted. The H-2D1 gene had a 15bp deletion where the sgRNA targeted (5’ CCGTNGGGTCGTTCTGTTCCAAACCTCGGACTTGGGACCCGGGACGTCAGCGTCCCTGTGTCGGGAAGTGGAGGGGCCTGACCTCCCACGCGGGGTCACTCACCGCCCGCGCTCTGGTTGTAGTAACCNAGCAGGTTCCTCAGGCTCACTCGGAACCACTGCTCTTGGGCCTTGGNTTTCTGTGTTTCCCGCTCCCAATACTCCGGCCCCTCCTGCTCCATCCACGGCGCCCGCGGCTCATATCTCGGATTCTCCGCGTCGCTTTCGAAACGCACGAACTCCTTGTTGTCCACATAGCCAACAGAGATGTACCGGGGC--------------- CGGGACACGGCGGTCTCGAAATACCGCATCGAGTGTGGGCCTGGGGACGGCGCGCGGTGAGACCCCGACCTCCTCACCAAACCCCGGGCGGCTGCGCACGCCGGGAGGGGATCTGGGCGCGGGGCTCAGGTGGAGAAGGGGCGGAGGGTCCGNGGGGGCGACGA 3’).

### Preparation of Cell Suspension

Thymi, and spleens were mechanically dissociated in FACS buffer (0.5% BSA in PBS) or complete RPMI (10% FBS) to generate single-cell suspensions that were then passed through a 70μm filter. Intraepithelial lymphocytes (IELs) were isolated from the small intestine as previously described^70^. Briefly, small intestine was cut to 1cm pieces and washed with cold CMF. Tissue pieces were allowed to settle and CMF was poured off. The tissue was then digested with DTE solution for 30 min at 37C in a 50mL conical tube. Tissue pieces were centrifuged at 1,500rpm for 5 min at 4C. Supernatant was collected and centrifuged at 1,500rpm for 5min at 4C. Lymphocytes were isolated by percoll separation utilizing 40% and 80% percoll ^21^. The percoll solution was centrifuged at 2000rpm with no brake for 20 min at room temperature. Lymphocyte layer was then washed with PBS and collected. Splenocytes were then RBC lysed using ACK lysis buffer (0.15M NH4Cl, 1mM KHCO3, 0.1mM Na2EDTA) for 5 minutes at room temperature.

### Staining for Flow Cytometry

Thymi, spleens, and IELs were stained in 2.4G2 supernatant for 30 minutes at 4°C with the following antibodies: (BD Biosciences) CD4 (RM4-4), CD8α (53-6.7), CD5 (53-7.3), PLZF (R17-809), (Biolegend) TCRβ (H57-597), CD8β (YTS156.7.7), B220 (RA3-6B2), Vα3.2 (RR3-16), CD45.2 (104), T-bet (4B10),(Invitrogen) CD8β (H35-17.2), CD24 (M1/69), integrin α4β7 (DATK32), CD69 (H1.2F3) (Tonbo) CD44 (IM7), and CD45.1 (A20). Cells were then washed in PBS and stained in Ghost Dye Violet 510 as described above. For intracellular staining, cells were fixed and permeabilized using the eBioscience FoxP3/ Transcription Factor Staining Buffer Set (ThermoFisher) according to manufacturer’s instructions. Biotinylated peptide-MHC monomers were obtained from the NIH Tetramer Facility (Atlanta, GA). Tetramers were assembled by conjugating the biotin-labeled monomers with PE-labeled streptavidin (Agilent, #PJRS27-1) according to NIH Tetramer Facility protocols. Cell numbers were calculated using AccuCheck Counting Beads for count and pipetting accuracy (Life Technologies #PCB100) according to manufacturer’s instructions. All antibodies were from BD Biosciences, Biolegend, Invitrogen, or Tonbo Biosciences. Samples were processed using a Fortessa X20 (BD Biosciences) and analyzed using FlowJo software. For defining CD8α+CD8βlow thymocyte population, we adjusted the gate based on wild type controls samples analyzed in parallel to experimental samples, such that 5-6% of the wild type CD8SP cells were CD8βlow. This gate was then applied to our experimental samples (as shown in Fig. 1E). Similarly, the CD8α+CD8βlow gate in the IEL population was placed based on wild type samples run in parallel and then applied to experimental samples (Supplementary Fig. 1C).

### Tetramer Enrichment

Single-cell suspensions of thymi and spleens were generated as described above. Cells were incubated in 50nM Datsatinib (_Sigma_ Aldrich, CDS023389-25MG) for 30 minutes at 37°C and then stained with tetramer in 2.4G2 supernatant for 1 hour at room temperature. After staining, cells were washed and incubated with Anti-PE MicroBeads (Miltenyi Biotec, #130-048-801) in MACS buffer (0.5% BSA) for 30 minutes at 4°C. Cells were then positively enriched for tetramer+ T cells using a magnetic column (Miltenyi Biotec.) according to manufacturer’s instructions and washed before extracellular staining.

### Bone Marrow Dendritic Cell Culture *In Vitro* Stimulation

Bone marrow cells were harvested as described above and RBC lysed using ACK lysis buffer (0.15M NH4Cl, 1mM KHCO3, 0.1mM Na2EDTA) for 5 minutes at room temperature. Bone marrow cells were resuspended in cRPMI and seeded at 5 × 10^6^ cells per milliliter in 24 well plates. Cells were supplemented with GM-CSF (Peprotech, #315-03-20UG) until day 4 and adhering cells were harvested on day 6 using EDTA. 6 × 10^5 CD^11c+MHC-II+ bone marrow cells per milliliter were seeded in 24 well plates. Preselection QFL thymocytes were generated by crossing QFL TCR transgenic mice onto a non-selecting, MHC-I deficient background (β2M-/-). Thymic single-cell suspensions were generated as described above. Thymocytes were resuspended in cRPMI and seeded at 4 × 10^6^ cells per milliliter.

### Generation of DC 2.4 MHC I and β2m Kos

DC 2.4 cells were obtained from UC Berkeley’s Cell Culture Facility. DC 2.4 cells were transduced using the lentiCas9-Blast vector (Addgene plasmid # 52962; http://n2t.net/addgene:52962; RRID: Addgene_52962) and selected with 10 ug/mL of Blasticidin (AG Scientific #3513-03-9). Guide-RNA sequences targeting mouse β2m, Qa-1^b^, H2-K^b^, and H2-D^b^ were selected using CRISPick (portals.broadinstitute.org/gppx/crispick/public), and sequences are as follows: β2m: TCACGCCACCCACCGGAGAA, Qa-1^b^: TACTACAATCAGAGTAACGA, H2-K^b^: GTACATGGAAGTCGGCTACG, H2-D^b^: AGATGTACCGGGGCTCCTCG. Guide-RNAs were stably transduced in SpCas9-expressing DC 2.4 cells using the LentiGuide-Neo vector and selected with 500 ug/mL of G418 (InvivoGen #108321-42-2). LentiGuide-Neo was a gift from Caroline Goujon (Addgene plasmid # 139449; http://n2t.net/addgene:139449; RRID: Addgene_139449). DC 2.4 cells that were negative for surface β2m, Qa-1^b^, Qa-1^b^ × H2-D^b^, Qa-1^b^ × H2-K^b^, and Qa-1^b^ × H2-D^b^ × H2-K^b^ were independently sorted out on BD FACSAria Fusion (BD Biosciences).

### *In Vitro* Stimulation with DC2.4 cells

WT, Qa1^b^KO, Qa1^b^K^b^D^b^KO, Qa1^b^K^b^KO, Qa1^b^D^b^KO, and β2mKO DC2.4 cells were plated in 48 Well Cell Culture Plates (Corning, Cat. No:3548) at 3×10^5^ cells per well. Cells were left to settle for 1 hour at 37°C 5% CO_2_. DC2.4 cells were then treated with 5ng/ml Recombinant Mouse IFN-γ (Biolegend, Cat. No:575302) for 24 hours, then Recombinant Mouse IFN-γ was washed off. Single-cell suspensions of QFLTgβ2mKO thymi were prepared as described above. Thymocytes were overlaid at 1×10^5^ cells per well and cultured at 37°C 5% CO_2_ for 24 hours, then harvested for flow cytometric analyses.

### Thymic Tissue Slice Cultures

Thymic lobes from wild type, Qa1^b^KO and β2mKO mice were gently isolated and any connective tissue was removed. Lobes were embedded into 4% agarose with a low melting point (GTG-NuSieve Agarose, Lonza) and sectioned into 400-500mm slices using a vibratome (VT1000S, Leica). Thymic slices were overlaid onto 0.4mm transwell inserts (Corning, Cat. No.: 353090) in 6 well tissue culture plates with enough cRPMI under the insert to reach the slices. Pre-selection QFL (2.5×10^5^) thymocytes were overlaid onto each slice and allowed to migrate for 3 hours, after which excess thymocytes were removed by gently washing with PBS. Slices were cultured at 37°C 5% CO_2 until_ harvested for analysis. For flow cytometry, thymic slices were mechanically dissociated into single-cell suspensions prior to staining.

### Bone Marrow Chimeras

Host mice were depleted of NK cells by I.P. injecting anti-NK1.1 (PK136, Leinco Technologies, #N123) at 100ug/100uL every 24Hrs for two days, for a total of 200ug of depleting antibody. Mice were irradiated in two doses of 600 rads (total of 1,200rads), with a resting period of 16hrs between doses. Mice were maintained on antibiotic water (Trimethoprim / Sulfamethoxazole) 4 weeks following irradiation. Bone marrow was harvested from the femur of donor mice using standard techniques. Red blood cells were lysed using ACK lysis buffer (0.15M NH4Cl, 1mM KHCO3, 0.1mM Na2EDTA) for 5 minutes at room temperature. Cells were depleted of CD4+ T cells by staining with CD4 PE-conjugated antibody (RM4-4) for 20 minutes at 4°C and then with Anti-PE MicroBeads (Miltenyi Biotec.) as described above. The labeled cells were washed, resuspended in MACS buffer, and then passed through a magnetic column (Miltenyi Biotec.). Flow-through (CD4-depleted bone marrow cells) was washed, resuspended at (4×10^6^ cells) in 100uL of PBS and i.v. injected into recipient mice. Bone marrow chimeras were analyzed 8-11 weeks following reconstitution.

### CFSE labeling

Cells were resuspended in 5uM CFSE proliferation dye (ThermoFisher #C34554) and incubated at 37°C for 9 minutes. Cells were then washed by addition of pre-warmed cRPMI while vortexing. Cells were then washed again with pre-warmed PBS and resuspended at the desired concentration.

### *In Vitro* Stimulation with Splenocytes

Single cell-suspensions of splenocytes were generated and RBC lysed as described above. Antigen presenting cells (APCs) were prepared by depleting splenocytes of CD4, CD8, and NK1.1 expressing cells using a magnetic column as described above. APCs were seeded at 4×10^5^ cells per well in a 48 well plates. Responding cells were splenocytes from QFLTg mice that were depleted of CD4, NK1.1, B220, and CD19 expressing cells using a magnetic column as described above. Cells were then CFSE (ThermoFisher #C34554) labeled as described above and seeded at 1×10^5^ cells per well.

### Generation of RAG2KO Neonatal Chimera

Donor thymocytes and splenocytes were isolated as described above. Single cell suspensions were depleted of CD4+ T cells, B cells and NK cells by staining with CD4 (RM4-4), CD19 (1D3), and NK1.1(PK136) PE-conjugated antibody for 20 minutes at 4°C and then with Anti-PE MicroBeads (Miltenyi Biotec) as described above. The labeled cells were washed, resuspended in cRPMI buffer, and then passed through a magnetic column (Miltenyi Biotec). Flow-through (CD4, CD19, NK1.1 depleted cells) was washed, resuspended at (1×10^6^ cells) in 100uL of PBS and intrahepatically injected into 4–6-day old Rag2KO recipients. Recipient mice were analyzed 14-15 weeks post-injection.

## Figure Legends

**Supplemental Figure 1:**
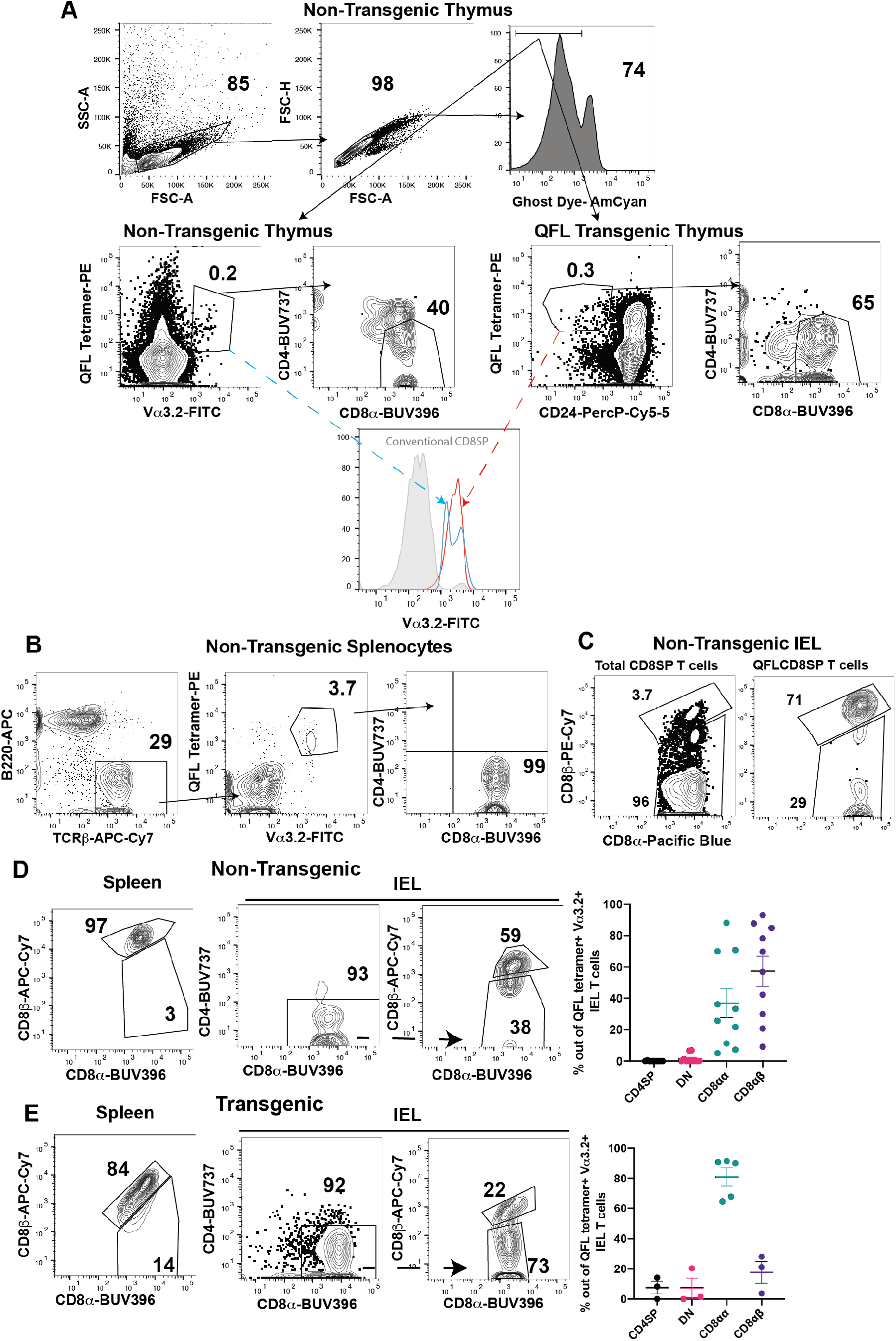
Gating Strategy utilized to identify QFL T cells in distinct tissues. **A)** Representative gating strategy employed to identify QFL CD8SP T cells in non-transgenic (tetramer enriched) and QFLTg thymocytes. Note that virtually all QFL tetramer^+^CD24^-^ from QFLTg mice express Vα3.2 (lower histogram). **B)** Representative plots showing the gating strategy to identify QFL CD8SP T cells in non-transgenic (tetramer enriched) and QFLTg spleen and IEL compartment of the small intestine. Example shown from Spleen of non-transgenic mice. **C)** Representative flow plots illustrating gates identifying CD8αβ and CD8α^+^βlow populations within CD8SP T cells in the IEL compartment **(Left)** Total CD8SP T cells (Gated: B220^-^TCRβ^+^CD8α^+^CD4^-^) **(Right)** QFL CD8SP T cells (Gated: B220^-^TCRβ^+^QFL tetramer^+^Vα3.2^+^CD8α^+^CD4^-^) **D-E) (Left)** Representative plots of CD8α and CD8β on QFL CD8SP splenocytes **(Middle)** Representative plots CD4, CD8α and CD8β of QFL T cells in the SI IEL compartment. (**Right)** Compiled data showing the % of indicated populations in QFL IEL compartment (Gated: TCRβ^+^QFL tetramer^+^Vα3.2^+^) **(D)** (n=10) and QFLTg **(E)** (n=5)mice. CD4SP (CD4^+^CD8α^-^) (Black dots), DN (CD4-CD8α^-^) (Magenta dots), CD8αα(CD4^-^CD8α^+^CD8β^-^) (Teal dots), CD8αβ (CD4^-^CD8α^+^CD8β^+^) (Purple dots). Comparisons that are not statistically significant are not marked by a symbol.

**Supplemental Figure 2:**
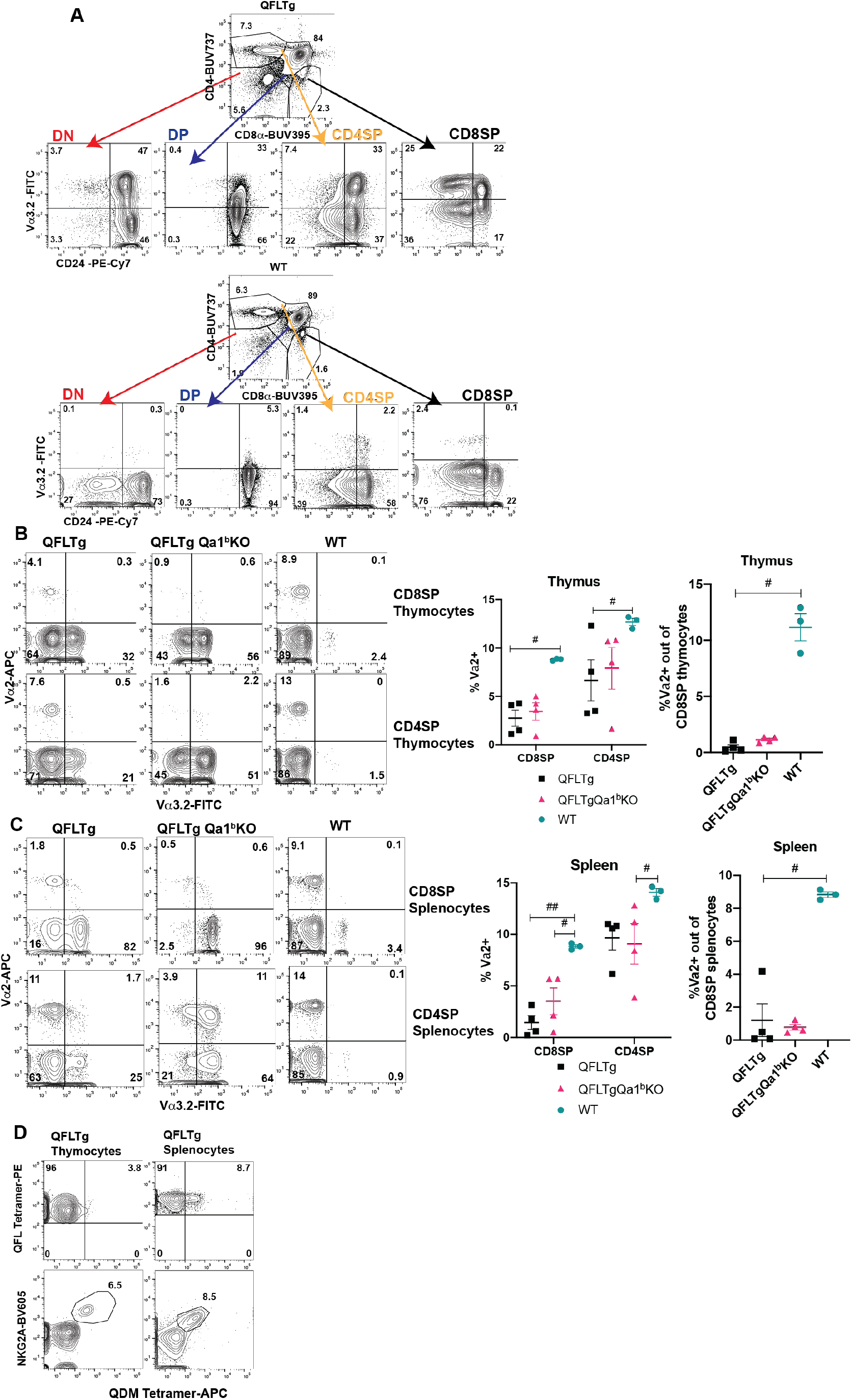
Characterization of QFL TCR transgenic mice. **A)** Representative flow plots of CD4 and CD8α expression on thymocytes from QFLTg mice (Gated: Live) or Vα3.2 and CD24 expression of indicated thymocyte populations. Data from wild type mice is show below for comparison. **B)** Representative flow plots of Vα2 and Vα3.2 expression in CD8SP (Gated: CD8α^+^CD4^-^CD24^-^) and CD4SP (Gated: CD8α^-^ CD4^+^CD24^-^) thymocytes from QFLTg, QFLTgQa1^b^KO and WT thymi. Compiled data of Vα2 expression is shown on the right of the plots. Compiled data of Vα2 expression on QFL CD8SP (Gated: QFL tetramer^+^CD24^-^CD8α^+^CD4^-^) thymocytes shown on the far right. **C)** Representative flow plots of Vα2 and Vα3.2 expression in CD8SP and CD4SP splenocytes from QFLTg, QFLTgQa1^b^KO, and WT spleens (Gated: TCRβ^+^B220^-^CD8α^+^CD4^-^ (CD8SP) or TCRβ^+^B220^-^CD8α^+^CD4^-^ (CD4SP)). Compiled data of Vα2 expression is shown on the right of the plots. Compiled data of Vα2 expression on QFL CD8SP (Gated: TCRβ^+^B220^-^QFL tetramer^+^CD8α^+^CD4^-^) splenocytes shown on the far right. Each dot represents an individual mouse. **D)** Representative flow plots of **(Top)** QFL tetramer and QDM tetramer and **(Bottom)** NKG2A and QDM tetramer on QFLTg thymocytes (Gated: QFL tetramer^+^CD24^-^) and QFLTg splenocytes (Gated: TCRβ^+^B220^-^QFL tetramer^+^). Error bars= Standard error of mean. Statistical analysis: One way ANOVA followed by Tukey’s multiple comparison test. # P value, <0.05, ## P value <0.005, ### P value<0.0005, #### P value<0.00001. Comparisons that are not statistically significant are not marked by a symbol.

**Supplemental Figure 3:**
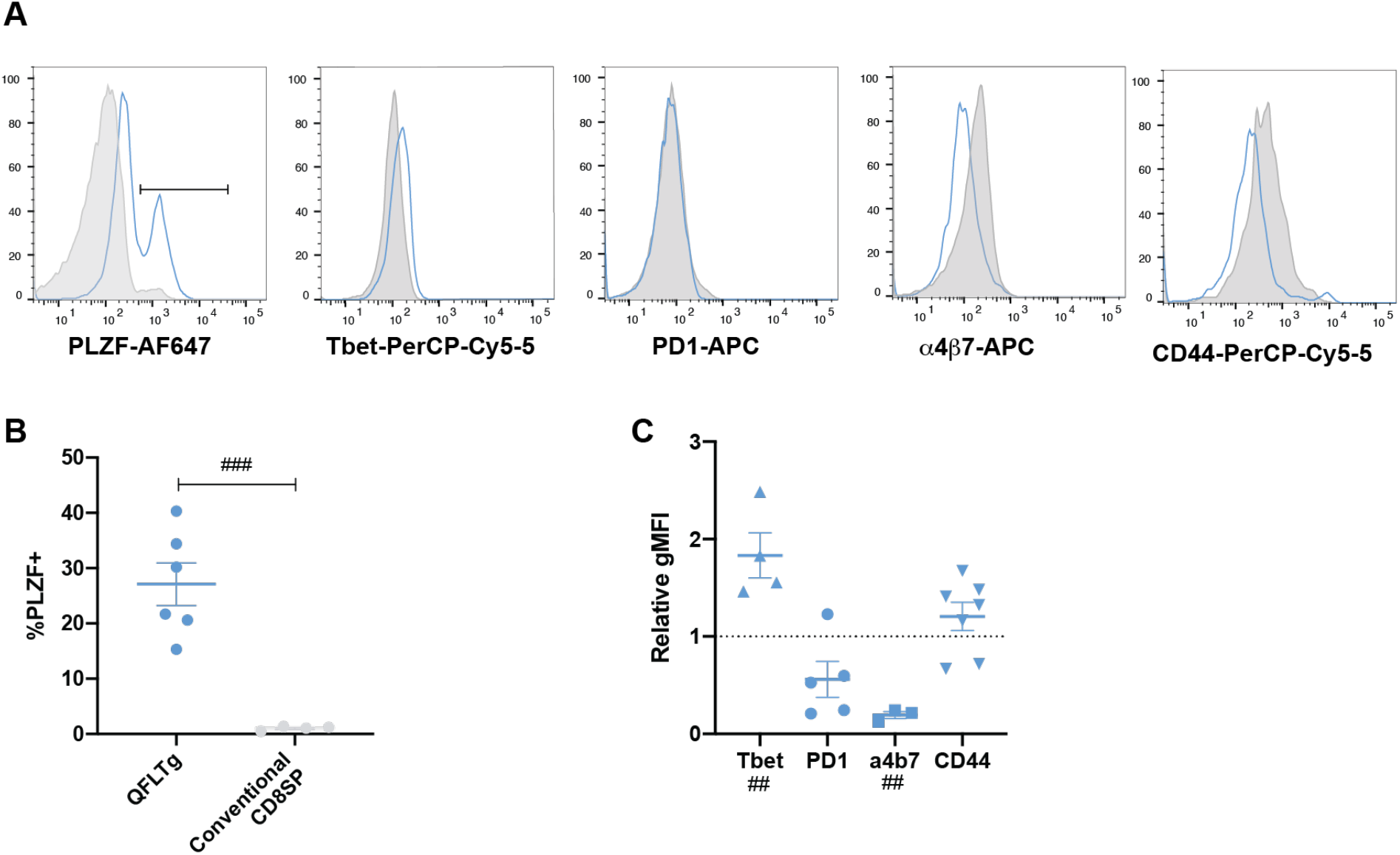
Characterization of QFL CD8SP thymocytes. **A)** Representative histograms of PLZF (n=6), Tbet (n=4), PD1 (n=5), α4β7(n=4) and CD44 (n=5) in QFL CD8SP thymocytes (Light Blue curve/dots) (Gated: QFL tetramer^+^CD24^-^ CD8α+CD4^-^). Conventional CD8SP T cells (Grey histogram/dots) (Gated: TCRβ^+^ CD8α^+^CD4^-^) are shown for comparison. **B)** Percentage of PLZF^+^ out of QFL CD8SP thymocytes or conventional CD8SP thymocytes. **C)** Quantification of Tbet, PD1, α4β7 and CD44 are expressed as a ratio of gMFI on QFL CD8SP thymocytes over conventional CD8SP thymocytes. Error bars= Standard error of mean. Statistical analysis: student’s T test **(B)** # P value, <0.05, ## P value <0.005, ### P value<0.0005, #### P value<0.00001One way ANOVA followed by Tukey’s multiple comparison test **(C)** # P value, <0.05, ## P value <0.005, ### P value<0.0005, #### P value<0.00001. Comparisons that are not statistically significant are not marked by a symbol.

**Supplemental Figure 4:**
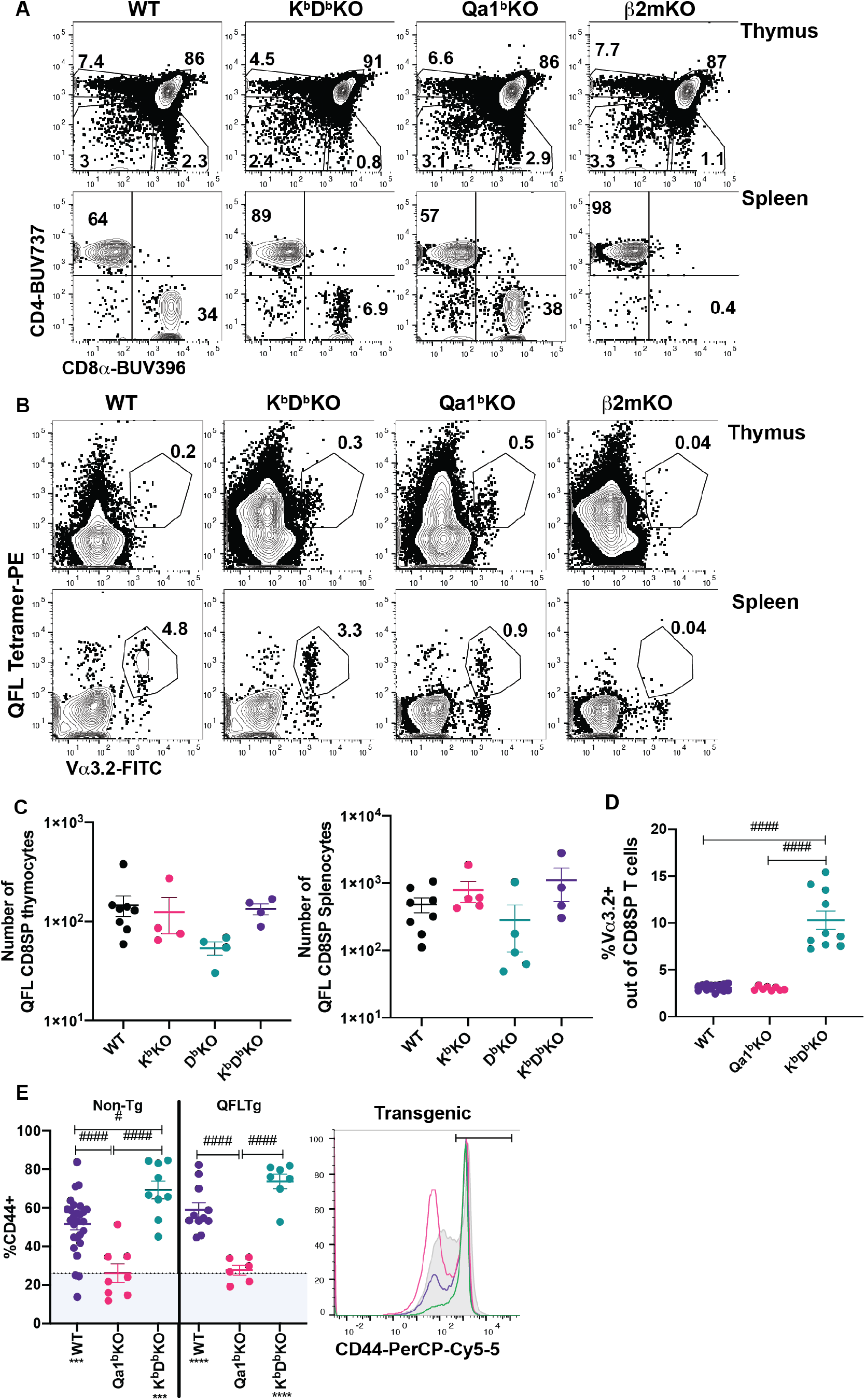
Phenotype of K^b^D^b^KO and Qa1^b^KO thymus and spleen. **A)** Representative flow plots of CD4 and CD8α expression on unenriched WT, K^b^D^b^KO, Qa1^b^KO and β2mKO thymocytes (Gated: Live cells) and splenocytes (Gated: TCRβ^+^). **B)** Representative flow plots of QFL tetramer and Vα3.2 expression on tetramer enriched thymocytes (Gated: Live) and splenocytes (Gated: TCRβ^+^) of B6, K^b^D^b^KO, Qa1^b^KO and β2mKO mice. **C)** Number of QFL T cells in thymi (tetramer enriched and gated: QFL tetramer^+^Vα3.2^+^CD8α^+^CD4^-^) or spleen (tetramer enriched and gated: TCRβ^+^QFL tetramer^+^Vα3.2^+^ CD8α^+^CD4^-^) of non-transgenic mice of the indicated genotype. (Thymus WT n=9, K^b^KO=6, D^b^KO=6 K^b^D^b^KO n=5) (Spleen WT n=8, K^b^KO=5, D^b^KO=5 K^b^D^b^KO n=4). **D)** Frequency of Vα3.2+ cells out of CD8SP splenocytes (Gated: TCRβ^+^B220^-^ CD8α^+^CD4^-^) from the indicated mouse strains (n=10 for all conditions). **E)** Representative histogram and compiled data of CD44 expression on QFL CD8SP T cells (Gated: TCRβ^+^QFL tetramer^+^Vα3.2^+^CD8α^+^) in Non-Transgenic (WT n=32, Qa1^b^KO n=8 and K^b^D^b^KO n=9) **(Left of the line)** and QFL Transgenic(WT n=11, Qa1^b^KO n=6 and K^b^D^b^KO n=7)**(Right of the line)**: WT (Purple dots), Qa1^b^KO (Magenta dots) and K^b^D^b^KO spleens. Dotted line represents the average value for conventional CD8SP (Gated: TCRβ^+^CD8α^+^) (22%). <u>Error</u> bars= Standard error of mean. Statistical analyses: One way ANOVA comparing samples to conventional CD8 T cells (shown below sample name). * P value <0.05, ** P value <0.005 *** P value<0.0005. One way ANOVA followed by Tukey’s multiple comparison test. # P value, <0.05, ## P value <0.005, ### P value<0.0005, #### P value<0.00001 Comparisons that are not statistically significant are not marked by a symbol.

**Supplemental Figure 5:**
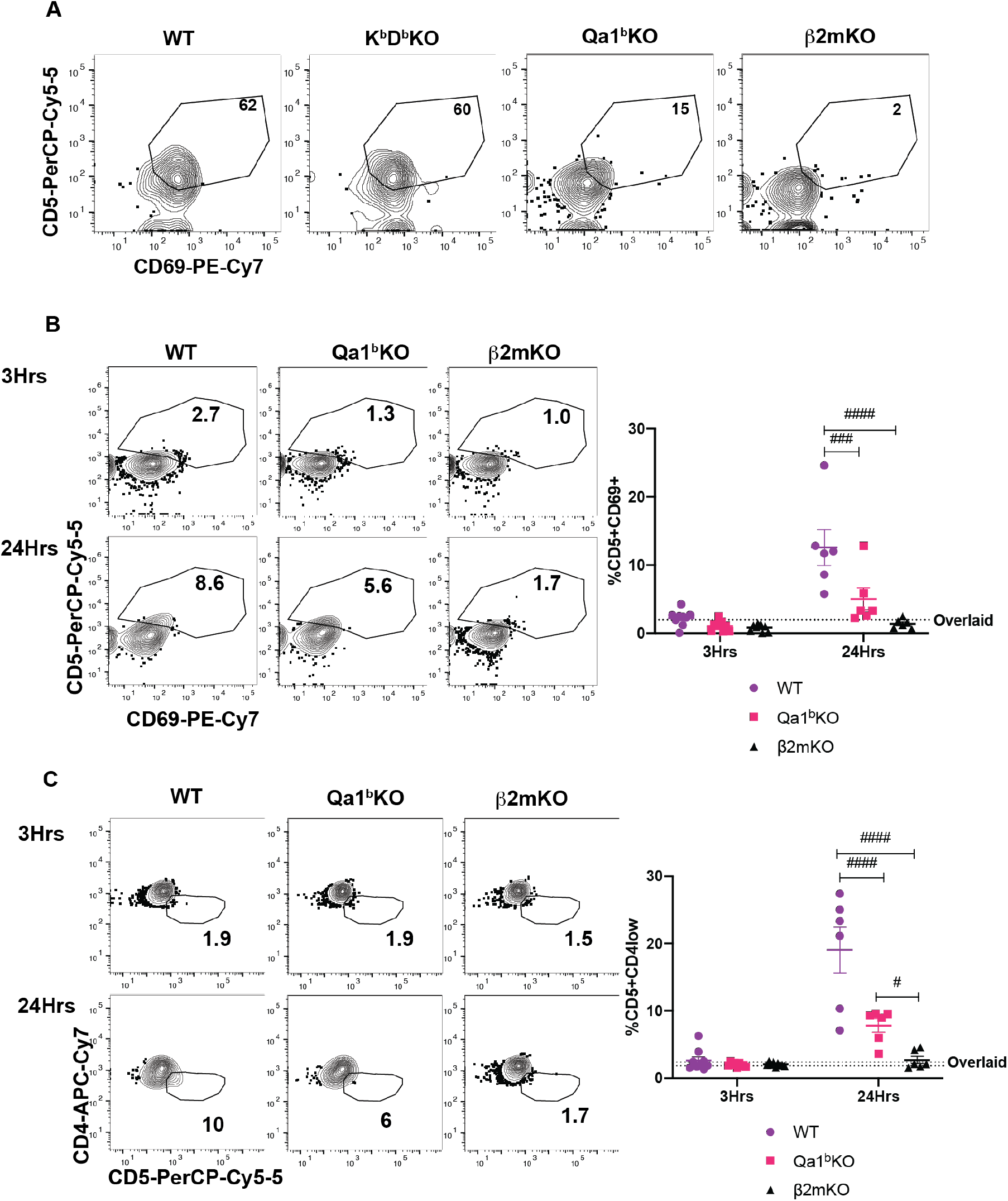
QFL thymocyte stimulation *in-vitro*. **A)** Representative flow cytometry plots of CD5 and CD69 expression on preQFLTg DP thymocytes (Gated: QFL tetramer^+^Vα3.2^+^CD4^+^CD8α^+^) after 24 hours of co-culture with Bone Marrow Derived Dendritic cells (BMDC) from the indicated mouse strains. **B-C)** Pre-selection QFL thymocytes (from QFLTg β2mKO mice) were overlaid onto thymic tissue slices from the indicated mouse strains. Representative flow cytometry plots of **(B)** CD5 and CD69 expression or **(C)** CD5 and CD4 expression on QFL DP thymocytes (Gated: QFL tetramer^+^Vα3.2^+^CD4^+^CD8α^+^) after 3 (n=5 for all conditions) and 24 (n=5 for all conditions) hours of co-culture. Dot plots show compiled data of two experiments, with each dot representing a sample from an individual thymic slice. Error bars= Standard error of mean. Statistical analysis: Two-way ANOVA followed by Tukey’s multiple comparison test. For comparisons across displayed samples (shown above dots) # P value, <0.05, ## P value <0.005, ### P value<0.0005, #### P value<0.00001 Comparisons that are not statistically significant are not marked by a symbol.

**Supplemental Figure 6:**
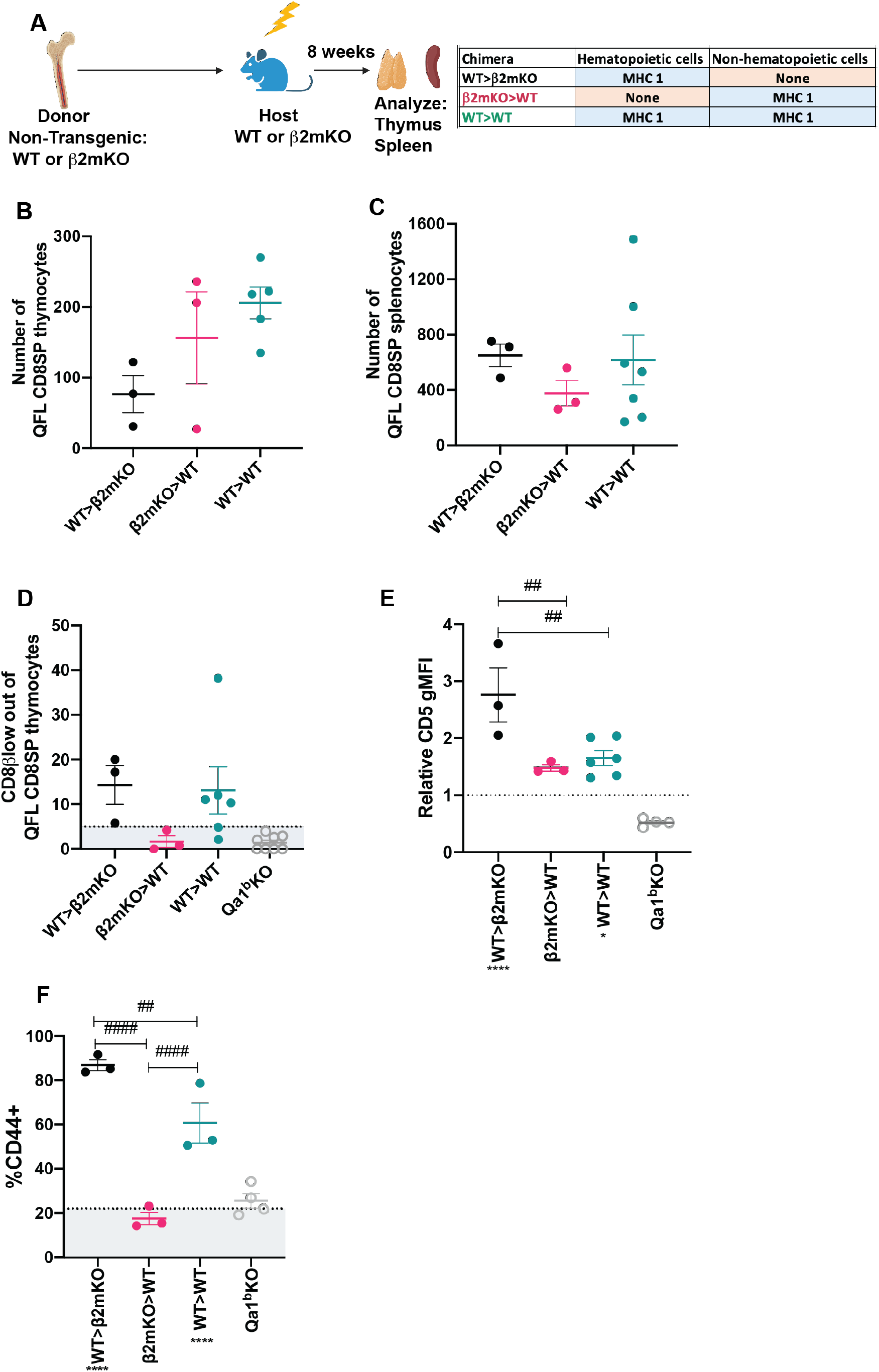
Requirement for hematopoietic cells versus non-hematopoietic cell MHC I expression in QFL T cell development in non-transgenic mice. **A)** Diagram of experimental design. Non-transgenic WT or β2mKO mice were used as bone marrow donors to reconstitute irradiated β2mKO or wild type hosts in order to restrict MHC I expression to hematopoietic or non-hematopoietic cells. **B-C)** Absolute numbers of QFL CD8SP T cells in **(B)** thymus (tetramer enriched and gated: QFL tetramer^+^Vα3.2^+^CD8α^+^) and **(C)** spleens (tetramer enriched and gated: TCRβ^+^QFL tetramer^+^Vα3.2^+^CD8α^+^) from the indicated chimeric mice(WT>β2mKO n=3, β2mKO>WT n=3, WT>WT n=5). **D)** Downregulation of CD8β of QFL CD8SP thymocytes of the indicated chimeric mice. Dotted line represents the average for conventional CD8SP (Gated: CD8α^+^CD4-) from unenriched non-transgenic thymi (5.2%) (WT>β2mKO n=10, β2mKO>WT n=10, WT>WT n=5). **E)** CD5 expression on QFL CD8SP thymocytes of the indicated non-transgenic chimeric mice and Qa1^b^KO mice. Graph shows gMFI of CD5 expression of QFL thymocytes normalized to the gMFI of conventional CD8SP thymocytes from wild type mice analyzed in the same experiment (WT>β2mKO n=10, β2mKO>WT n=10, WT>WT n=5). **F)** Quantification of CD44 expression of QFL CD8SP T cells (Gated: TCRβ^+^QFL tetramer^+^Vα3.2^+^CD8α^+^CD4^-^) from tetramer enriched non-transgenic splenocytes from: WT>β2mKO (Black dots), β2mKO>WT (Magenta dots), WT>WT (Teal dots) chimeric spleens and Qa1^b^KO (Light blue dots) spleens. Dotted line represents conventional CD8SP (Gated: TCRβ^+^CD8α^+^) (22%-26%) (n=3 for all conditions). Error bars= Standard error of mean. Statistical analysis: Kruskal-Wallis test followed by Dunn’s test multiple comparison test (Panel B and C). For comparisons across displayed samples (shown above dots) # P value, <0.0332, ## P value <0.0021, ### P value<0.0002, #### P value<0.00001. One way ANOVA comparing sample to Conventional CD8SP. * P value <0.05, ** P value <0.005 *** P value<0.0005. One way ANOVA followed by Tukey’s multiple comparison test. # P value, <0.05, ## P value <0.005, ### P value<0.0005, #### P value<0.00001. Comparisons that are not statistically significant are not marked by a symbol.

